# Transcriptional control of parallel-acting pathways that remove discrete presynaptic proteins in remodeling neurons

**DOI:** 10.1101/2020.02.21.959700

**Authors:** Tyne W. Miller-Fleming, Andrea Cuentas-Condori, Laura Manning, Sierra Palumbos, Janet E. Richmond, David M. Miller

## Abstract

Synapses are actively dismantled to mediate circuit refinement, but the developmental pathways that regulate synaptic disassembly are largely unknown. We have previously shown that the epithelial sodium channel UNC-8 triggers an activity-dependent mechanism that drives the removal of presynaptic proteins liprin-*α*/SYD-2, Synaptobrevin/SNB-1, RAB-3 and Endophilin/UNC-57 in remodeling GABAergic neurons in *C. elegans* (Miller-Fleming et al., 2016). Here, we report that the transcription factor Iroquois/IRX-1 regulates UNC-8 expression as well as an additional pathway, independent of UNC-8, that functions in parallel to dismantle functional presynaptic terminals. We show that the additional IRX-1-regulated pathway is selectively required for the removal of the presynaptic proteins, Munc13/UNC-13 and ELKS, which normally mediate synaptic vesicle fusion and neurotransmitter release. Our findings are notable because they highlight the key role of transcriptional regulation in synapse elimination and reveal parallel-acting pathways that orchestrate synaptic disassembly by removing specific active zone proteins.

## INTRODUCTION

The nervous system is actively remodeled during development as new synapses are assembled and others are removed to refine functional circuits. In some cases, synaptic remodeling is limited to a specific developmental stage in which activity drives circuit plasticity. These “critical periods” are indicative of the necessary role of genetic programs that define these developmental windows for activity-induced remodeling. Thus, synaptic remodeling mechanisms are likely to depend on the combined effects of both transcriptionally-regulated and activity-dependent pathways (Hensch 2004; Kano and Watanabe 2019).

In the nematode, *C. elegans*, synapses in the GABAergic motor circuit are relocated by a stereotypical remodeling program during early larval development. Dorsal D (DD) GABAergic motor neurons are generated in the embryo and initially synapse with ventral body muscles (Figure 1A). During the first larval stage, presynaptic domains are removed from ventral DD processes and then reassembled in the dorsal nerve cord (Figure 1B) (White, Albertson, and Anness 1978). Postembryonic Ventral D (VD) neurons are born during this early larval period and synapse exclusively with ventral muscles (Figure 1B). In the resultant mature circuit, alternating GABAergic output to dorsal (DD) versus ventral (VD) muscles is required for sinusoidal movement (White et al. 1976, 1986).

**Figure 1:**
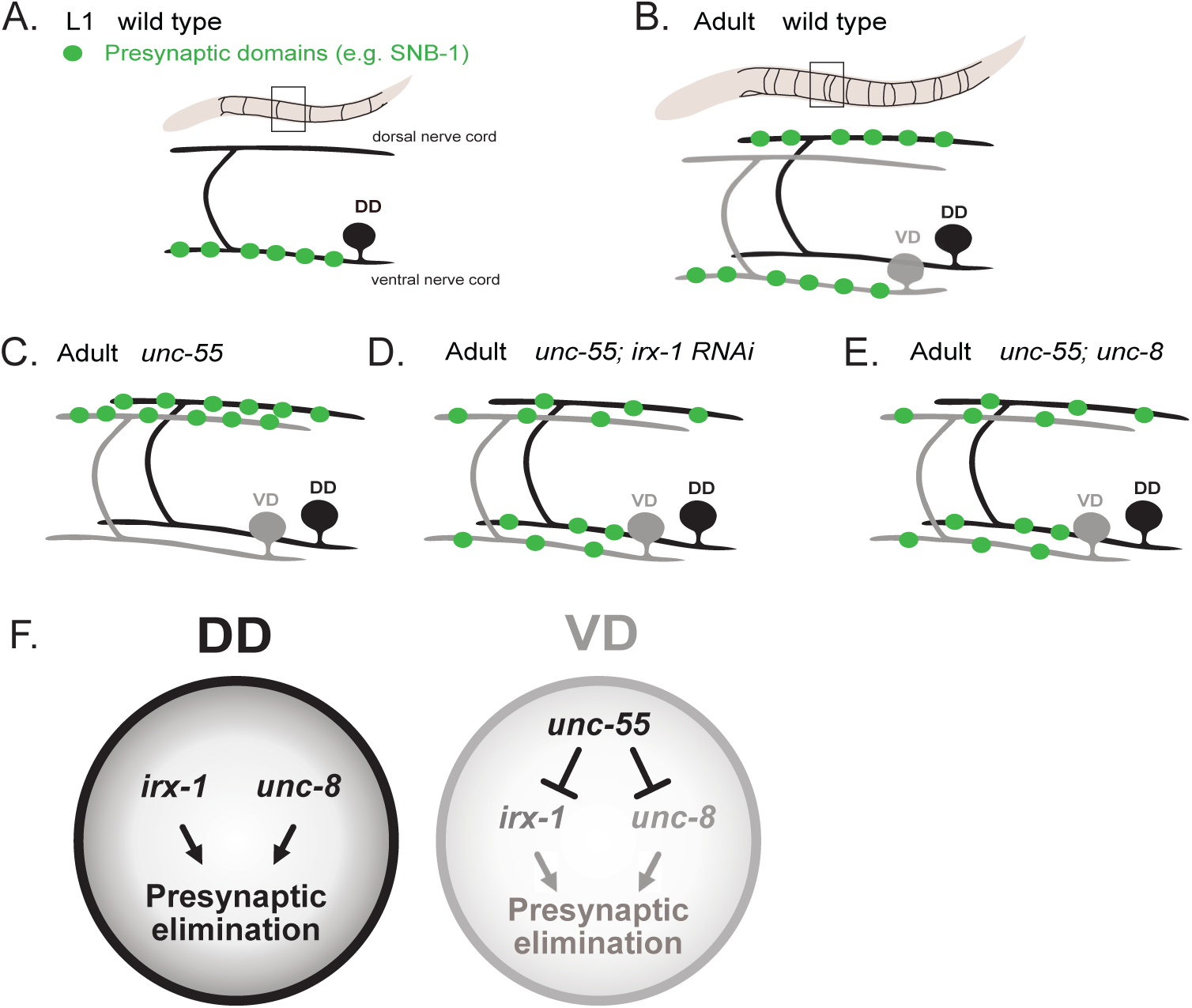
A transcriptional program regulates GABAergic neuron synaptic remodeling. **A.** DD motor neurons innervate ventral muscles in the early L1 stage. GFP-tagged synaptobrevin (SNB-1::GFP) marks GABAergic presynaptic domains. **B.** DD synapses are relocated to the dorsal nerve cord during early larval development as postembryonic VD class GABAergic motor neurons are generated to synapse with ventral muscles. These DD and VD connections are maintained in the adult motor circuit. **C.** In *unc-55* mutants, both DD and VD presynaptic domains are relocated to the dorsal nerve cord. **D.** RNAi knock down of the Iroquois family homeodomain transcription factor, IRX-1, antagonizes GABAergic neuron synaptic remodeling in *unc-55* mutants. **E.** Mutations that disable the DEG/ENaC cation channel, UNC-8, impair the removal of DD and VD GABAergic presynaptic domains in *unc-55* mutants. **F.** *irx-1* and *unc-8* are normally expressed in DD neurons to drive disassembly of the presynaptic apparatus. The COUP-TF transcription factor, UNC-55, blocks expression of *irx-1* and *unc-8* in VD neurons to prevent synapse elimination.

The COUP-TF transcription factor, UNC-55, is selectively expressed in VD neurons to prevent synaptic remodeling; in *unc-55* mutants, VD neurons initially synapse with ventral muscles but then mimic the native DD remodeling program by relocating presynaptic domains to the dorsal nerve cord (Figure 1C) (Shan et al. 2005; Zhou and Walthall 1998). The idea that UNC-55 blocks expression of genes that normally drive synaptic remodeling is supported by the finding that forced expression of UNC-55 in DD neurons antagonizes the native remodeling program (Shan et al. 2005). In earlier work, we exploited the ectopic synaptic remodeling phenotype of *unc-55* mutants in cell-specific profiling experiments to identify UNC-55 targets. An RNAi screen detected a subset of *unc-55*-regulated genes that are required for synaptic remodeling. For example, RNAi knockdown of the Iroquois homeodomain transcription factor, IRX-1, reduced removal of GABAergic presynaptic domains from the ventral nerve cord in *unc-55* mutants (Figure 1D) (Petersen et al. 2011). Similarly, a loss-of-function allele of the DEG/ENaC cation channel subunit gene, *unc-8*, also antagonized synaptic remodeling in *unc-55* mutants (Figure 1E). Additional experiments confirmed that both the IRX-1 transcription factor and UNC-8 cation channel normally promote the native DD remodeling program (Figure 1F) (Miller-Fleming et al. 2016).

DEG/ENaC proteins function as cation channels and we have previously shown that UNC-8 gates sodium influx (Matthewman et al. 2016; Wang et al. 2013). UNC-8 promotes presynaptic disassembly in a pathway that depends on intracellular calcium and neural activity (Miller-Fleming et al. 2016). Here we show that DEG/ENaC/UNC-8 is transcriptionally-regulated by Iroquois/IRX-1 to remove the presynaptic components liprin-*α*/SYD-2, Synaptobrevin/SNB-1, RAB-3 and Endophilin/UNC-57. Our findings indicate that these presynaptic proteins are also disassembled by a separate Iroquois/IRX-1-dependent pathway that functions in parallel to DEG/ENaC/UNC-8. Thus, remodeling of GABAergic synapses depends on the combined effects of neural activity (UNC-8) and developmentally-regulated transcription (IRX-1). Finally, we show that the active zone proteins Munc13/UNC-13 and ELKS-1 are exclusively removed by Iroquois/IRX-1, but not UNC-8. Thus, our work suggests that synaptic disassembly in the GABAergic circuit is orchestrated by parallel-acting mechanisms that selectively target molecularly distinct components of the presynaptic apparatus during the remodeling period.

## RESULTS

### The homeodomain transcription factor, Iroquois/IRX-1, drives DEG/ENaC/UNC-8 expression in remodeling GABAergic neurons

In previous work, we used gene expression profiling and an RNAi screen to identify protein-encoding genes that promote presynaptic disassembly in remodeling GABAergic neurons (Petersen et al. 2011). Because these studies determined that two of these proteins, the homeobox transcription factor IRX-1/Iroquois and the DEG/ENaC ion channel subunit UNC-8 are both involved in removing the presynaptic vesicle SNARE protein, synaptobrevin/SNB-1 (Miller-Fleming et al. 2016; Petersen et al. 2011), we decided to test the hypothesis that Iroquois/IRX-1 regulates DEG/ENaC/UNC-8 expression.

We have previously shown that GFP reporter lines for the *irx-1* and *unc-8* genes are highly expressed in DD motor neurons but not in VD motor neurons in a wild-type background (Miller-Fleming et al. 2016; Petersen et al. 2011) (Figure 1). Here, we used single-molecule Fluorescent In-Situ Hybridization (smFISH) to detect *unc-8* transcripts in remodeling DD neurons and to determine if the *irx-1* gene is necessary for *unc-8* expression. DD neurons in which *irx-1* is targeted by cell-specific RNAi (csRNAi) (See Methods) showed significantly fewer *unc-8* transcripts in comparison to the wildtype (Figure 2A-B), suggesting that Iroquois/IRX-1 is required for DEG/ENaC/UNC-8 expression in DD neurons. Because forced expression of Iroquois/IRX-1 in VD motor neurons is sufficient to drive the elimination of VD presynaptic terminals (Petersen et al. 2011), we next asked if Iroquois/IRX-1 over-expression (*irx-1-*OE) could also induce *unc-8* expression in VD neurons. smFISH quantification confirmed that *unc-8* transcripts are elevated in *irx-1-OE* VD neurons in comparison to wildtype (Figure 2C-D). Together, these results demonstrate that the transcription factor, Iroquois/IRX-1, is both necessary and sufficient for DEG/ENaC/UNC-8 expression in remodeling GABA neurons (Figure 2E).

**Figure 2:**
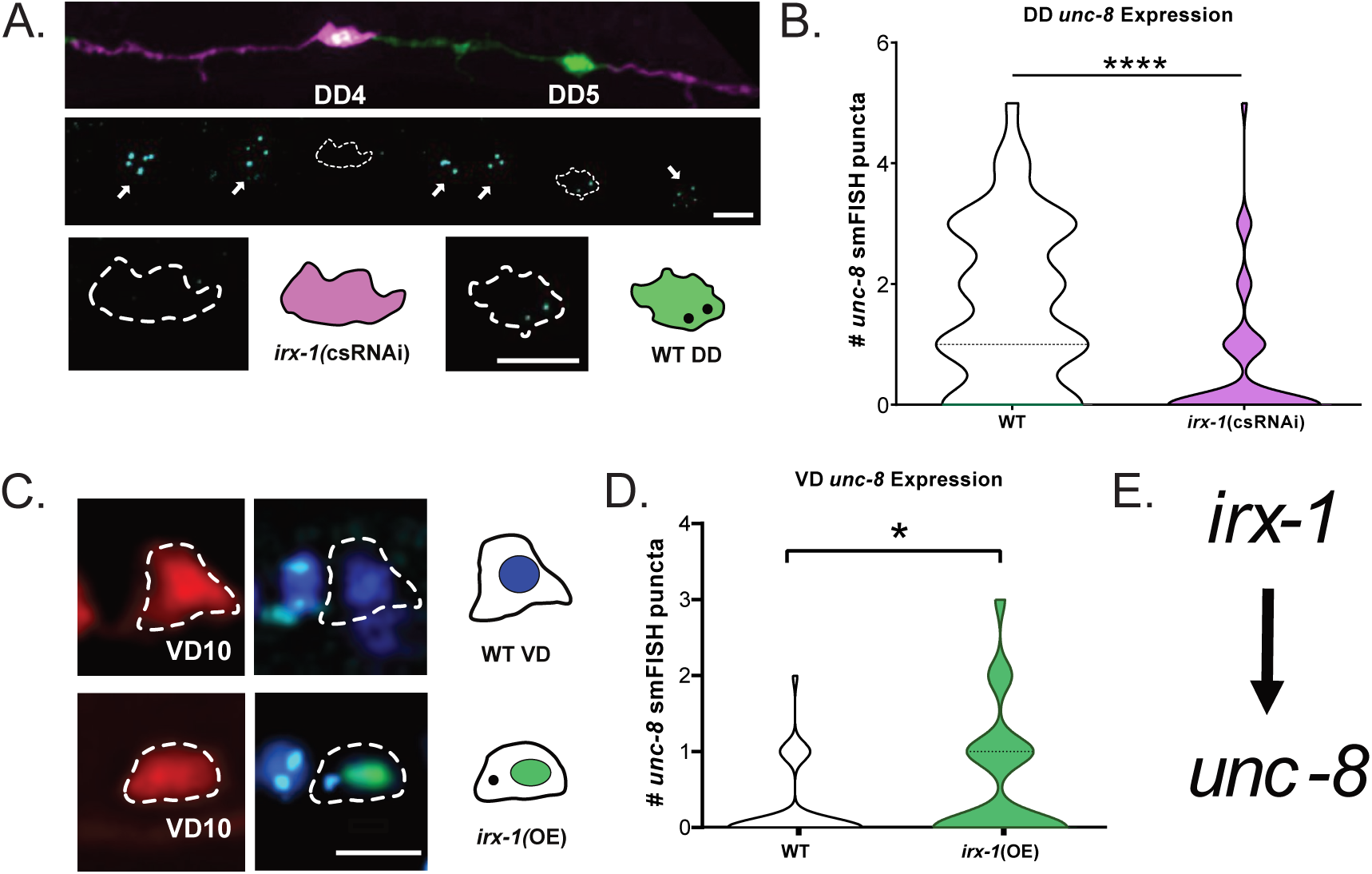
IRX-1 drives expression of UNC-8/DEG/ENaC to eliminate GABAergic presynaptic terminals. **A.** *irx-1(csRNAi)* (cell-specific RNAi) blocks *unc-8* expression in DD neurons. (Top) Mosaic expression of *irx-1(csRNAi)* (red) in DD4 vs adjacent by wild-type DD5 neuron (green) in the L1 ventral nerve cord. (Middle) *unc-8* smFISH puncta. Dashed lines demarcate DD cell soma and arrows denote *unc-8* smFISH puncta in adjacent cholinergic motor neurons. (Bottom) Dashed outlines and graphical representations depict *irx-1(csRNAi)-*marked DD4 neuron and *unc-8* smFISH puncta in wild-type (WT) DD5 neuron. Scale bar = 5μm. **B.** Quantification of *unc-8* smFISH puncta in DD neurons. Violin plots for *unc-8* smFISH puncta in wild-type (WT) (white) (n = 49) vs *irx-1(csRNAi*) (magenta) (n = 50) in L1 stage DD motor neurons. Dashed line represents median. Mann-Whitney test was used to determine significance. P<.0001 **C.** Over-expression of IRX-1 in VD neurons promotes *unc-8* expression. Representative images of VD neurons (VD10) in WT (top) vs *irx-1(OE)* (bottom). Dashed lines denote cell soma marked with mCherry (left) and labeled with DAPI (dark blue) and *unc-8* smFISH probe (light blue puncta) (right). Note IRX-1::GFP labels VD10 nucleus in *irx-1(OE)* (bottom right). Scale bar = 5μm. **D.** Quantification of *unc-8* smFISH puncta in VD motor neurons. Violin plots for *unc-8* smFISH puncta in WT (white) (n = 40) and *irx-1(OE)* (green) (n = 50) in L3 stage VD neurons. Dashed line represents median. Mann-Whitney test was used to determine significance. P = 0.0184 **E.** Working model: IRX-1 promotes UNC-8 expression in GABAergic motor neurons.

### Iroquois/IRX-1 drives a DEG/ENaC/UNC-8-dependent mechanism of presynaptic disassembly as well as a separate parallel-acting pathway that does not require UNC-8 for synaptic removal

We have previously shown that Iroquois/IRX-1 promotes the removal of Synaptobrevin/SNB-1::GFP from ventral presynaptic domains in remodeling GABAergic neurons (Figure 1E) (Petersen et al. 2011). If Iroquois/IRX-1 activates UNC-8 expression as predicted by our smFISH results (Figure 2), then Iroquois/IRX-1 should also drive removal of the additional presynaptic markers liprin-*α*/SYD-2, endophilin/UNC-57 and RAB-3/GTPase which depends on UNC-8 function (Miller-Fleming et al. 2016). To test for this possibility, we exploited *unc-55* mutants in which the VD GABAergic presynaptic domains are eliminated due to ectopic activation of the native DD remodeling program (Zhou and Walthall 1998) (Figure 1C). In this paradigm, removal of ventral GABAergic synapses in *unc-55* mutants is prevented by mutations that disable the pro-remodeling program. For example, ventral SNB-1::GFP puncta are eliminated in *unc-55* mutant animals but a significant fraction is retained in *unc-55; unc-8* double mutants (Figures 1E and 3A). This result confirms our earlier finding that UNC-8 function is required for the efficient removal of presynaptic SNB-1::GFP in remodeling GABAergic neurons (Miller-Fleming et al. 2016). Similarly, RNAi knockdown of *irx-1* prevents the elimination of ventral SNB-1::GFP puncta in *unc-55* worms (Figures 1D & 3A) (Petersen et al. 2011). Additional experiments showed that ablation of either *unc-8* or *irx-1* also blocks the removal of SYD-2::GFP and RAB-3::mCherry (Figure 3B-C) in the ventral nerve cord of *unc-55* mutants. Since Iroquois/IRX-1 induces *unc-8* expression (Figure 2), these results are consistent with the hypothesis that Iroquois/IRX-1 drives an UNC-8-dependent mechanism to remove presynaptic terminals in GABAergic neurons.

**Figure 3:**
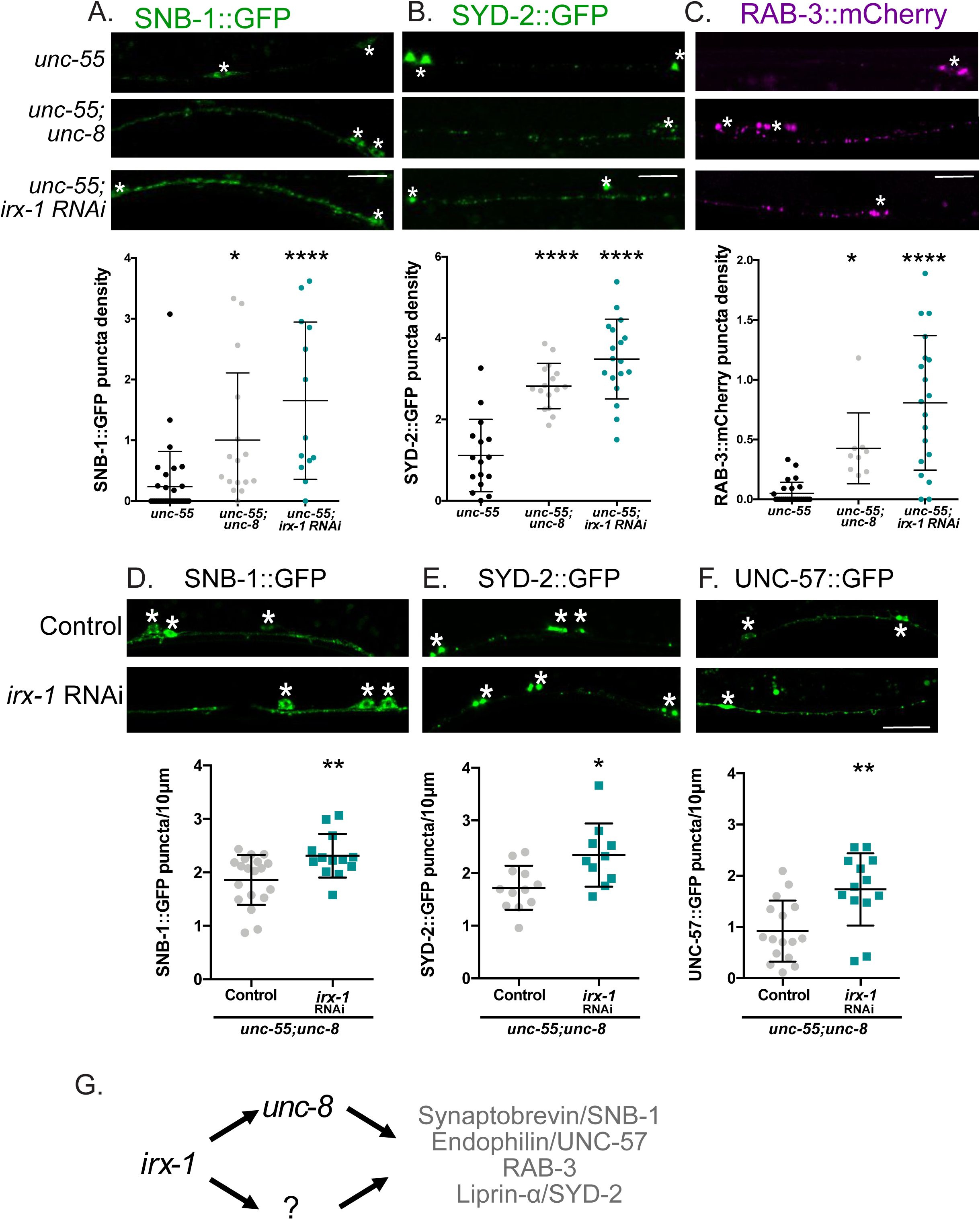
IRX-1 dismantles the GABAergic presynaptic apparatus via UNC-8-dependent mechanisms and with a parallel acting pathway that does not require UNC-8. **A.** (Top) Representative images of SNB-1::GFP-labeled GABA neuron synapses in the ventral nerve cord. (Bottom) SNB-1::GFP puncta density (puncta/10 μm) quantified for each genotype: *unc-55; unc-8* (1.0 ± 1.1) and *unc-55; irx-1(RNAi)* (1.65 ± 1.3) show more ventral SNB-1::GFP-marked puncta than *unc-55* (0.23 ± 0.57). Data are mean ± Standard Deviation (SD). *** = p<0.001, **** = p<0.0001. Kruskal-Wallis test with multiple comparison because *unc-55* dataset is not normally distributed. N > 12 animals. **B.** (Top) Representative images of SYD-2::GFP-labeled GABA neuron synapses in the ventral nerve cord. (Bottom) SYD-2::GFP puncta density (puncta/10 μm) quantified for each genotype: *unc-55; unc-8* (2.82 ± 0.6) and *unc-55; irx-1(RNAi)* (3.48 ± 0.9) show more ventral SYD-2::GFP-marked puncta than *unc-55* (1.1 ± 0.9). Data are mean ± SD. **** = p<0.0001. One-Way ANOVA with multiple comparison because all datasets are normally distributed. N > 15 animals. **C.** (Top) Representative images of RAB-3::mCherry-labeled GABA neuron synapses in the ventral nerve cord. (Bottom) RAB-3::mCherry puncta density (puncta/10 μm) quantified for each genotype: *unc-55; unc-8* (0.43 ± 0.3) and *unc-55; irx-1(RNAi)* (0.81 ± 0.6) show more ventral RAB-3::mCherry-marked puncta than *unc-55* (0.05 ± 0.1). Data is Mean ± SD. *** = p<0.001, **** = p<0.0001. Kruskal-Wallis test with multiple comparison because *unc-55* data is not normally distributed. N > 8 animals. **D.** *irx-1*-RNAi-treated *unc-55; unc-8* worms show more ventral SNB-1::GFP puncta/10 μm (2.31 ± 0.4) than *unc-55; unc-8* controls (1.86 ± 0.5). Data is Mean ± SD. N >12. Unpaired T-test. **p = 0.0086. **E.** *irx-1* RNAi-treated *unc-55; unc-8* worms show more SYD-2::GFP puncta/10 μm (2.34 ± 0.6) than *unc-55; unc-8* controls (1.72 ± 0.4). Data is Mean ± SD. N > 9. Unpaired T-test. * p = 0.01. **F.** *irx-1* RNAi-treated *unc-55; unc-8* worms show more UNC-57::GFP puncta/10 μm (1.73 ± 0.7) than *unc-55;unc-8* controls (0.92 ± 0.6). Data is Mean ± SD. N > 12. Unpaired T-test. ** p = 0.0023. **G.** Working model: IRX-1 drives removal of SNB-1, UNC-57, RAB-3 and SYD-2 from ventral GABA neuron presynaptic domains via an *unc-8*-dependent mechanism and with a separate parallel-acting pathway that does not require *unc-8*. All images of L4 stage larva, anterior to left. Asterisks mark GABAergic motor neuron cell soma. Scale bars =10 μm.

Because *irx-1* encodes a transcription factor, we reasoned that Iroquois/IRX-1 might also regulate other targets in addition to the *unc-8* gene in the GABA neuron synaptic remodeling pathway. This idea is consistent with our finding that wild-type levels of presynaptic components are not fully restored in *unc-55; unc-8* double mutants thus pointing to an additional parallel acting mechanism for synaptic disassembly (Miller-Fleming et al. 2016). We reasoned that if Iroquois/IRX-1 regulates a downstream target that functions in tandem with UNC-8, then genetic ablation of *irx-1* in an *unc-55; unc-8* double mutant should enhance the retention of ventral presynaptic markers in comparison to *unc-55; unc-8* mutants. For this test, we used feeding RNAi for global knockdown of *irx-1* because the *irx-1* null allele is lethal (Petersen et al. 2011). We counted GFP puncta for the presynaptic proteins SNB-1::GFP, SYD-2::GFP and UNC-57::GFP in *irx-1*-RNAi-treated *unc-55; unc-8* mutants. These experiments showed that RNAi knockdown of *irx-1* increases the number of ventral SNB-1::GFP, SYD-2::GFP and UNC-57::GFP puncta (Figure 3D-F) in *unc-55; unc-8* double mutant animals. Together, these results demonstrate that Iroquois/IRX-1 drives an additional genetic pathway, independent of UNC-8, that eliminates presynaptic terminals in remodeling GABAergic neurons (Figure 3G).

### Iroquois/IRX-1, but not DEG/ENaC/UNC-8, removes fusion-competent synaptic vesicles in remodeling GABAergic neurons

The retention of presynaptic markers (e.g., SNB-1::GFP, SYD-2::GFP) (Figure 3D-F) in ventral GABA motor neuron processes of *unc-55; unc-8* double mutants suggested that a reconstituted presynaptic apparatus in these animals might also restore synaptic release. We previously used electron microscopy (EM) to confirm the presence of ventral cord GABAergic synapses in *unc-55; unc-8* adults (Figure 4A) (Miller-Fleming et al. 2016). In contrast, EM analysis did not detect GABAergic ventral cord synapses in *unc-55* animals as expected since both DD and VD motor neuron synapses are relocated to the dorsal nerve cord in *unc-55* mutants (Miller-Fleming et al. 2016; Walthall and Plunkett 1995). Since our assays with fluorescent presynaptic markers also showed that Iroquois/IRX-1 drives presynaptic disassembly (Figure 3), we used EM to ask if cell-specific RNAi (csRNAi) knockdown of *irx-1* would prevent the removal of ventral GABAergic synapses in an *unc-55* mutant. These experiments detected GABAergic presynaptic terminals in *unc-55; irx-1(csRNAi)* samples (Figure 4A) (7 synapses/613 sections, n = 3 animals), thus, confirming that IRX-1 is necessary for the removal of synapses in remodeling GABA neurons (See Methods). Together, these EM data are consistent with our findings that ventral puncta for fluorescently-labeled SNB-1, SYD-2, RAB-3 and UNC-57 are absent in *unc-55* mutants, but that knockdown of either *unc-8* or *irx-1* activity partially rescues these defects (Figure 3A-C).

**Figure 4:**
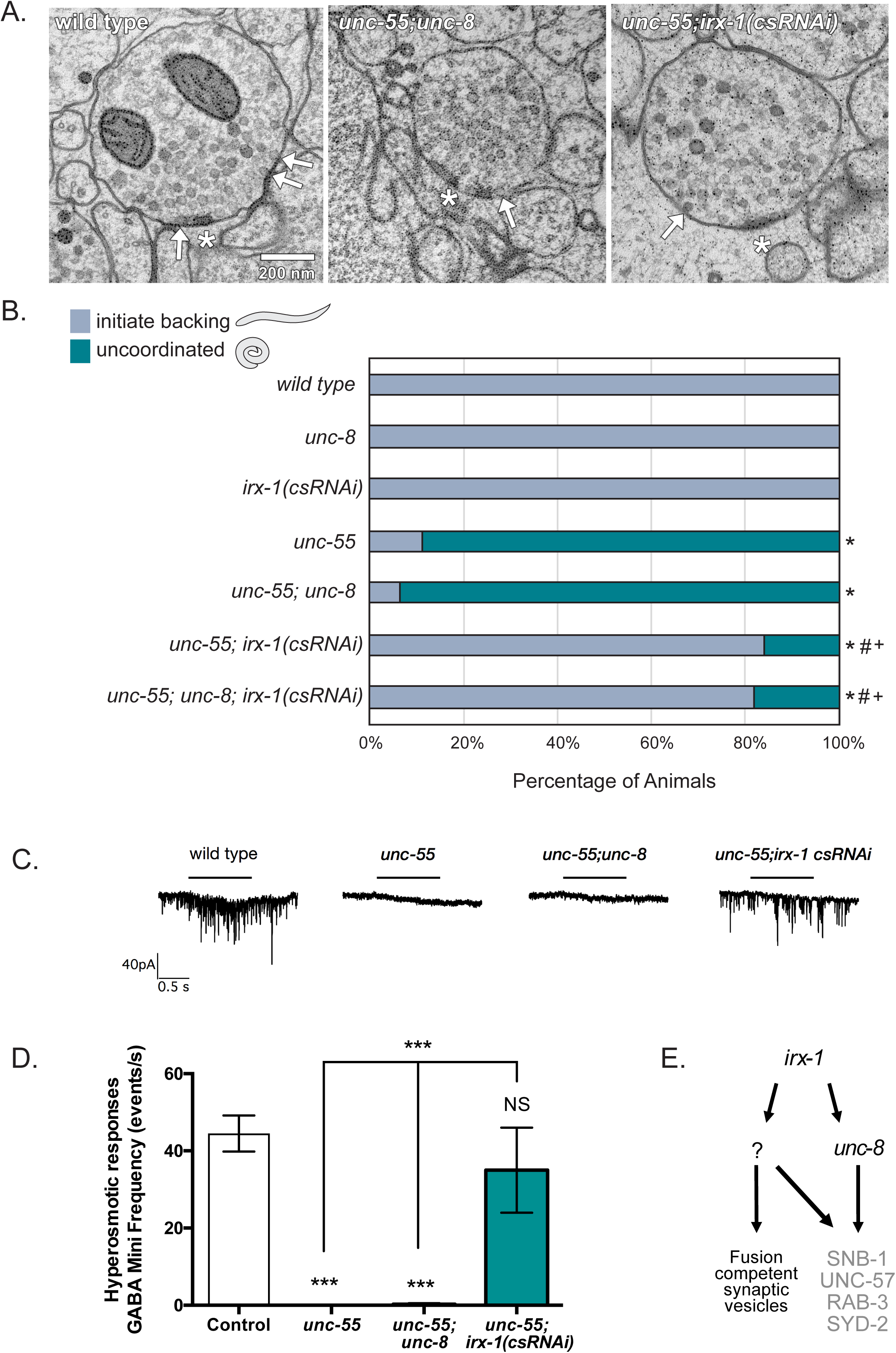
Genetic knockdown of *irx-1*, but not *unc-8*, restores ventral GABA synaptic function to *unc-55* mutants. **A.** Representative electron micrographs of GABA neuron synapses with ventral muscles for wild type, *unc-55; unc-8,* and *unc-55*; *irx-1(csRNAi).* Asterisks denote presynaptic density. Arrows point to docked synaptic vesicles. Scale bar = 200 nm. **B.** Animals were tapped on the head and scored for wild-type (gray) versus uncoordinated (teal) backward movement. *unc-55* and *unc-55; unc-8* animals coil ventrally with head tap (NS, not significant, Fisher’s Exact Test, n ≥ 100 animals per genotype) indicating that a loss-of-function *unc-8* mutation does not rescue backward locomotion in *unc-55* mutants. Cell-specific RNAi (*csRNAi*) knock-down of *irx-1* restores backward locomotion to *unc-55* animals and this effect is not enhanced in *unc-55; unc-8; irx-1(csRNAi).* * denotes significantly different from wildtype with p < 0.002. # denotes significantly different from *unc-55* with p < 0.002. + denotes significantly different from *unc-55; unc-8* with p < 0.002. See Figure 3S1 for complete list of statistical comparisons. **C.** Representative traces showing that ventral mini-iPSCs are detected for wild-type and *unc-55; irx-1(csRNAi)* animals, but not for *unc-55* or *unc-55; unc-8* mutants prior to and after application of hyperosmotic saline. Horizontal lines denote hyperosmotic treatment. **D.** Hyperosmotic treatment evokes iPSCs that are not significantly (NS) different between wild type (57.3 ± 6.8) vs *unc-55; IRX-1 csRNAi* (50.7± 15). In contrast, GABA mini frequencies in *unc-55* (0 ± 0) and *unc-55; unc-8* animals (0 ±0.25) in response to hyperosmotic treatment were significantly diminished (N ≥ 3, data are mean ± SEM, One-Way ANOVA Bonferroni correction, *** p = 0.001). **E.** Working Model: *irx-1* drives expression of *unc-8* and also activates an additional pathway involving unknown downstream components that removes fusion-competent synaptic vesicles from remodeling GABAergic synapses.

Our EM results determined that genetic ablation of *unc-8* restores the ultrastructural morphology of some ventral GABAergic synapses to *unc-55* animals (Figure 4A), including the presence of presynaptic dense projections, synaptic vesicles and plasma-membrane docked vesicles (Miller-Fleming et al. 2016). As a first test of the possibility that these synapses are functional, we used a behavioral readout that depends on ventral GABAergic synapses. Ventral synapses for both DD and VD neurons are dismantled in *unc-55* mutants and reassembled in the dorsal nerve cord (Figure 1D). The resultant imbalance due to excess inhibitory GABAergic output to dorsal muscles versus excess excitatory cholinergic input to ventral muscles results in a striking behavioral phenotype in which *unc-55* animals coil ventrally when tapped on the head instead of initiating coordinated backward locomotion (Shan et al. 2005; Walthall and Plunkett 1995). We have shown that mutants that disable pro-remodeling genes (*e.g., unc-8, irx-1*) restore ventral GABAergic synapses to *unc-55* mutants (Figure 1C). If these restored synapses are functional, then the tapping assay should detect improved backward locomotion in double mutants with *unc-55*. In this assay, each animal is tapped gently on the head to induce backward movement. We determined, however, that *unc-55; unc-8* double mutant animals display severely defective backward locomotion that is not significantly different from that of *unc-55* mutants (Figures 4B & 4S1). This finding suggests that that ventral GABAergic synapses in *unc-55; unc-8* mutants are not functional. In contrast, *unc-55; irx-1(csRNAi)* animals (Figure 4B) show robust backward movement in comparison to *unc-55* mutants. This result suggests that GABAergic release is restored to ventral cord synapses by RNAi knockdown of *irx-1* and thus that Iroquois/IRX-1 is required for the removal of functional GABAergic synapses in *unc-55* mutants. (Figure 4B,C) (Petersen et al. 2011). Genetic ablation of *unc-8* activity in this genetic background (i.e., *unc-55; unc-8; irx-1(csRNAi)*) does not further enhance backward locomotion as predicted by our conclusion that residual ventral cord GABAerigic synapses in *unc-55; unc-8* double mutants are not functional (Figure 4B) and by our finding that Iroquois/IRX-1 regulates expression of the *unc-8* gene (Figure 2). To summarize, the results of the movement assay suggest that although ventral GABAergic synapses are visible by EM in both *unc-55; unc-8* and in *unc-55; irx-1(csRNAi)* animals, GABAergic release is selectively reactivated by knockdown of *irx-1,* but not by genetic removal of *unc-8*.

To investigate the mechanism of this differential effect on synaptic function, we quantified the number and distribution of docked synaptic vesicles in *unc-55; unc-8* and *unc-55; irx-1(csRNAi)* double mutants. EM analysis determined that *unc-55; unc-8* and *unc-55; irx-1(csRNAi)* animals show similar numbers of docked synaptic vesicles in ventral cord GABAergic synapses (Figure 4S2). This finding argues against a model in which synaptic dysfunction in *unc-55; unc-8* mutants can be attributed to the failure of synaptic vesicles to contact the presynaptic membrane prior to neurotransmitter release.

We next examined whether the docked synaptic vesicles in *unc-55; unc-8* animals are fusion-competent by recording iPSCs from ventral muscles using d-tubocurare (dTBC) to block cholinergic signaling. As previously reported, tonic release of ventral iPSCs was restored in *unc-55; irx-1(csRNAi)* animals, but this was not observed in *unc-55; unc-8* mutants despite the presence of organized clusters of fluorescent presynaptic proteins (Figure 3), electron dense active zones (Figure 4) and docked vesicles in both strains (Figure 4S2) (Miller-Fleming et al. 2016; Petersen et al. 2011). Previously, we determined that the postsynaptic GABA_A_ receptor UNC-49 was also properly localized and functional in *unc-55; unc-8* animals (Miller-Fleming et al. 2016), thus excluding the possibility that the absence of iPSCs in *unc-55; unc-8* animals is due to a postsynaptic defect. The lack of tonic release in *unc-55; unc-8* animals could be due to defective vesicle priming or downstream Ca*^2+^-*sensing. To determine whether the morphologically docked synaptic vesicles in *unc-55; unc-8* animals are primed, we measured iPSCs in response to hypertonic sucrose perfusion which is sufficient to induce neurotransmitter release from the primed synaptic vesicle pool (Figure 4C-D). As expected, hyperosmotic stimulation triggered robust iPSCs in wild-type and in *unc-55; irx-1(csRNAi)* animals, whereas ventral muscles in *unc-55* mutants were unresponsive. Hyperosmotic treatment of *unc-55; unc-8* animals also failed to trigger ventral GABA release suggesting that these animals are defective in synaptic vesicle priming (Figure 4C-D). To rule out the possibility of a general defect in synaptic vesicle fusion, we confirmed that preparations of *unc-55* and *unc-55;unc-8* mutants exhibited spontaneous cholinergic activity that could be abolished by dTBC (data not shown). Together, these data are consistent with the idea that synaptic vesicles at ventral GABAergic synapses in *unc-55; unc-8* animals are capable of docking with the presynaptic membrane, but are not fusion competent. Because hyperosmotic treatment evokes robust iPSCs in *unc-55; irx-1(csRNAi)* animals we further conclude that Iroquois/IRX-1 also functions in the remodeling program to activate an additional *unc-8*-independent pathway that disassembles synaptic components required for synaptic vesicle priming (Figure 4E).

Previous work has shown that synaptic vesicle docking at the presynaptic active zone depends on the vesicular GTPase protein RAB-3 and the RAB-3-interacting protein RIM1/UNC-10 (Gracheva et al. 2008). RAB-3 is largely absent from the ventral cord processes of GABAergic neurons of *unc-55* animals as a result of ectopic VD remodeling (Miller-Fleming et al. 2016; Thompson-Peer et al. 2012), but *unc-55; unc-8* animals show ventral RAB-3 puncta (Figure 3C) (Miller-Fleming et al. 2016). We determined that fluorescently-labeled RIM1/UNC-10 is also localized to ventral GABAergic synapses in *unc-55; unc-8* double mutants. Surprisingly, unlike other presynaptic markers, RIM-1/UNC-10::GFP does not remodel in *unc-55* mutants and is retained on the ventral side (Figure 4S3). Together, these results suggest that dual localization of both RAB-3 and RIM1/UNC-10 in ventral GABAergic synapses of *unc-55; unc-8* mutants could account for our EM observation of normal numbers of docked vesicles (Figure 4S2-3).

### Iroquois/IRX-1 removes synaptic vesicle priming protein UNC-13 and the RIM-binding protein ELKS-1 in remodeling GABAergic neurons

The cytosolic protein Munc13/UNC-13 functions as a conserved component of the presynaptic apparatus to mediate synaptic vesicle fusion (Richmond, Davis, and Jorgensen 1999; T. C. Südhof 2012; Weimer et al. 2006). Mammalian neurons express four UNC-13-related proteins whereas only two distinct UNC-13 proteins, a long (UNC-13L) and a short (UNC-13S) version, are expressed in *C. elegans*. Because synaptic vesicles appear docked, but are incapable of fusion in *unc-55; unc-8* mutants (Figure 4), we hypothesized that UNC-13 could be missing from ventral GABAergic synapses in these animals. To test this idea, we generated a strain expressing GFP-tagged UNC-13L protein in GABA neurons. We selected UNC-13L for this experiment because it co-localizes with UNC-10/RIM whereas the short isoform, UNC-13S, shows a diffuse distribution in GABAergic motor neurons (Figure 5S1A), as also reported for *C. elegans* cholinergic motor neurons (Hu, Tong, and Kaplan 2013). As previously observed for other presynaptic proteins (Figure 3) (Hallam and Jin 1998; Miller-Fleming et al. 2016; Petersen et al. 2011; Thompson-Peer et al. 2012), UNC-13L::GFP is restricted to the ventral nerve cord prior to DD remodeling in early L1 larvae, but is detectable post-remodeling in both the dorsal and ventral nerve cords in adults (Figure 5S1B-C). This finding indicates that UNC-13L::GFP remodels to DD presynaptic domains in the dorsal nerve cord and is also a component of ventral VD synapses in the adult. We quantified the number of UNC-13L::GFP puncta in the ventral nerve cord and determined that UNC-13L::GFP is largely removed in *unc-55* mutants (Figure 5A). In contrast to other presynaptic markers (e.g. SNB-1::GFP) (Figure 3A-C), UNC-13L::GFP is also eliminated from ventral GABAergic synapses in *unc-55; unc-8* mutants (Figure 5A). Thus, wild-type UNC-8 activity is not required for the removal of UNC-13L from remodeling GABAergic synapses. This finding suggests that although the ventral presynaptic active zone in *unc-55; unc-8* mutant GABAergic neurons appears normal by EM (Figure 4A and 4S2) (Miller-Fleming et al. 2016), UNC-13L is not localized at these terminals thus likely accounting for their synaptic vesicle fusion defect (Figure 4C-D).

**Figure 5:**
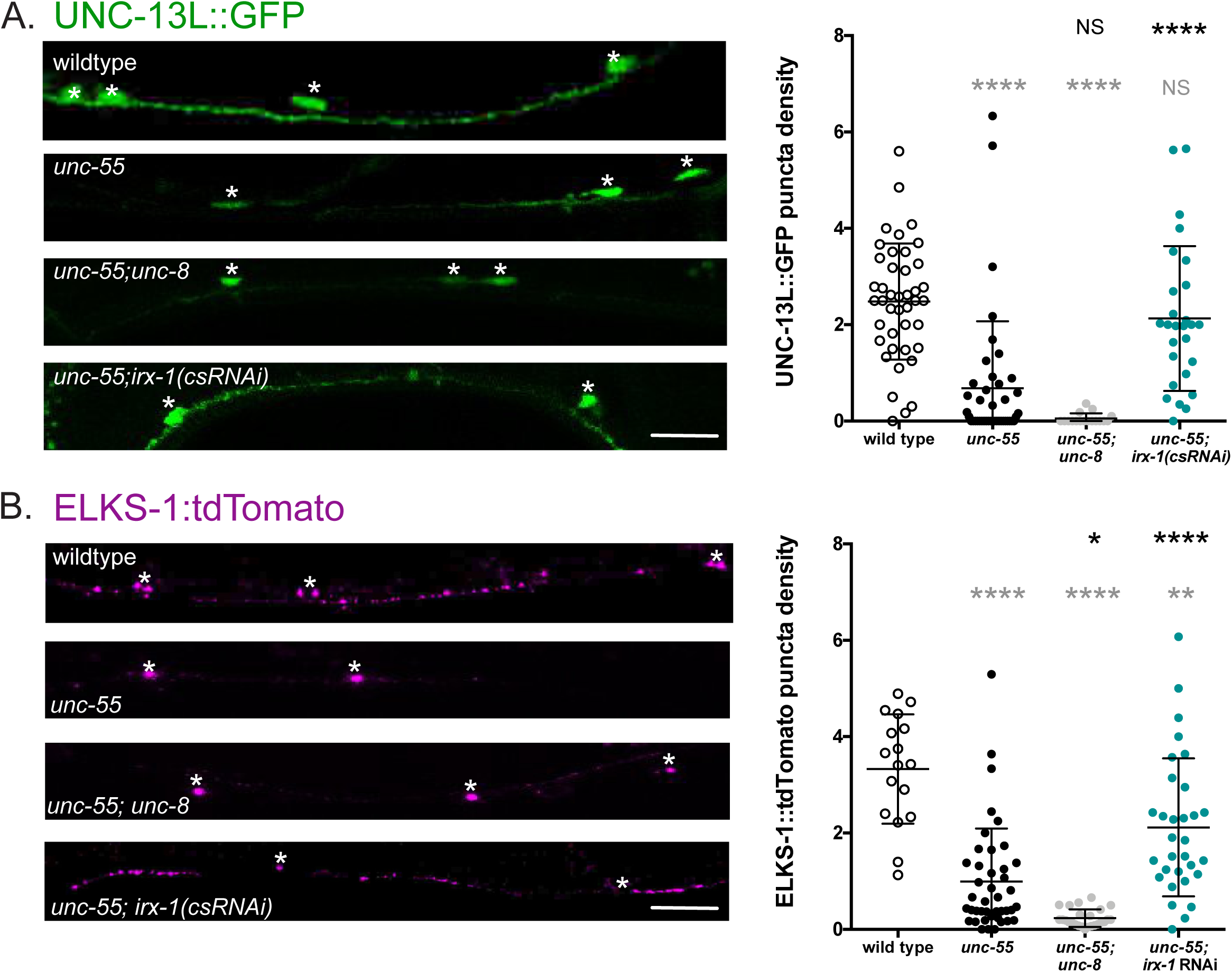
IRX-1, but not UNC-8, drives removal of Munc13/UNC-13 and ELKS-1 from the presynaptic domains of remodeling GABAergic neurons. **A.** (Left) Representative images of UNC-13L::GFP-labeled GABA neuron synapses in the ventral nerve cord. Asterisks denote cell soma. GABAergic UNC-13::GFP ventral puncta are missing in *unc-55* and in *unc-55; unc-8*, but are restored in *unc-55; irx-1(csRNAi)*. Scale bar = 10 μm. Images from L4 animals, anterior to left. (Right) Quantification of UNC-13L::GFP puncta at ventral GABAergic synapses of wildtype (2.48 ± 1.2), *unc-55* (0.68 ± 1.4), *unc-55; unc-8* (0.05 ± 0.1) and *unc-55; irx-1(csRNAi)* (2.19 ± 1.5). puncta density = puncta/10 μm. Data are mean ± SD. N > 16. One-Way ANOVA with Bonferroni correction. Gray asterisks denote comparisons vs wildtype. Black asterisks show comparisons vs *unc-55*. **** p < 0.0001. NS = not significant. **B.** (Left) Representative images of the presynaptic protein ELKS-1 indicate that *irx-1,* and not *unc-8,* is required for the removal of ELKS-1::tdTomato from ventral GABAergic synapses. Asterisks denote cell soma. Scale bar = 10 μm. Images from L4 animals. Anterior to left. (Right) Quantification of ventral GABAergic synapses labeled with ELKS-1::tdTomato in wildtype (3.3 ± 1.1), *unc-55* (0.99 ± 1.1), *unc-55; unc-8* (0.23 ± 0.2) and *unc-55; irx-1(RNAi)* (2.11 ± 1.4). puncta density = puncta/10 μm. Data are mean ± SD. N > 16. One-Way ANOVA with Bonferroni correction. Gray asterisks show comparisons vs wild-type. Black asterisks show comparisons vs *unc-55*. * p < 0.05; **** p < 0.0001. NS = not significant.

Since UNC-8 expression in VD neurons was shown to drive elimination of SNB-1::GFP (Miller-Fleming et al. 2016), we devised an experiment to test the idea that removal of UNC-13L is UNC-8-independent. We confirmed that forced expression of UNC-8 in VD neurons is sufficient to remove SNB-1::GFP from ventral GABAergic synapses, but does not displace UNC-13L::GFP (Figure 5S3). Together, these results show that UNC-8 function is neither necessary nor sufficient for UNC-13L removal from remodeling GABAergic synapses.

RNAi knock down of *irx-1* in *unc-55* mutants is sufficient to restore ventral GABAergic synaptic release (Figure 4C-D). Thus, we next asked if UNC-13L::GFP is retained in the ventral nerve cord of *unc-55; irx-1(csRNAi)* animals. Indeed, we found that ventral UNC-13L::GFP puncta are detectable in both wild-type and in *unc-55; irx-1(csRNAi)* animals (Figure 5A) thus indicating that Iroquois/IRX-1 function is required for the removal of UNC-13L from remodeling GABAergic synapses.

Based on previous work demonstrating that the RIM-binding protein ELKS-1 recruits the mammalian protein bMunc-13-2 to active zones and Drosophila ELKS homologue, Bruchpilot, recruits UNC-13L/Unc13A (Böhme et al. 2016; Kawabe et al. 2017), we next asked if ELKS-1 synaptic localization is differentially affected by the UNC-8 versus IRX-1-dependent synaptic remodeling pathways. We determined that expression of ELKS-1::tdTomato (Cherra and Jin 2016) in wild-type adult GABA neurons results in bright fluorescent puncta characteristic of DD synapses in the dorsal nerve cord and VD synapses in the ventral nerve cord (Figure 5S2). Ventral ELKS-1::tdTomato-labeled puncta are largely absent in *unc-55* mutants indicating that ELKS-1 is dismantled from the presynaptic domains of remodeling GABAergic motor neurons (Figure 5B). Ventral ELKS-1 puncta are also missing in *unc-55; unc-8* mutants. Thus, UNC-8 function is not required to remove ELKS-1 from the presynaptic domain (Figure 5B). In contrast, a substantial number of ELKS-1 puncta are retained in the ventral nerve cord of *unc-55; irx-1(csRNAi)* animals (Figure 5B). Therefore, our results are consistent with the idea that Iroquois/IRX-1 removes ELKS-1 from ventral synapses of remodeling GABAergic motor neurons in a genetic pathway that is independent of UNC-8 (Figure 7). To summarize, our results show that IRX-1/Iroquois drives the removal of multiple components of the presynaptic apparatus in remodeling GABAergic neurons including Synaptobrevin/SNB-1, RAB-3, liprin-*α*/SYD-2, Endophilin/UNC-57, UNC-13L and ELKS-1 (Figures 3 and 5). In contrast, DEG/ENaC/UNC-8 promotes the disassembly of Synaptobrevin/SNB-1, RAB-3, liprin-*α*/SYD-2 and Endophilin/UNC-57 (Figure 3) but is not required for the removal of Munc-13/UNC-13L and ELKS-1 from remodeling GABAergic presynaptic domains (Figures 5, 6 and 5S3). These findings are notable because they show that distinct genetic pathways can dismantle the presynaptic apparatus in remodeling GABAergic neurons by targeting specific active zone components.

**FIGURE 6.**
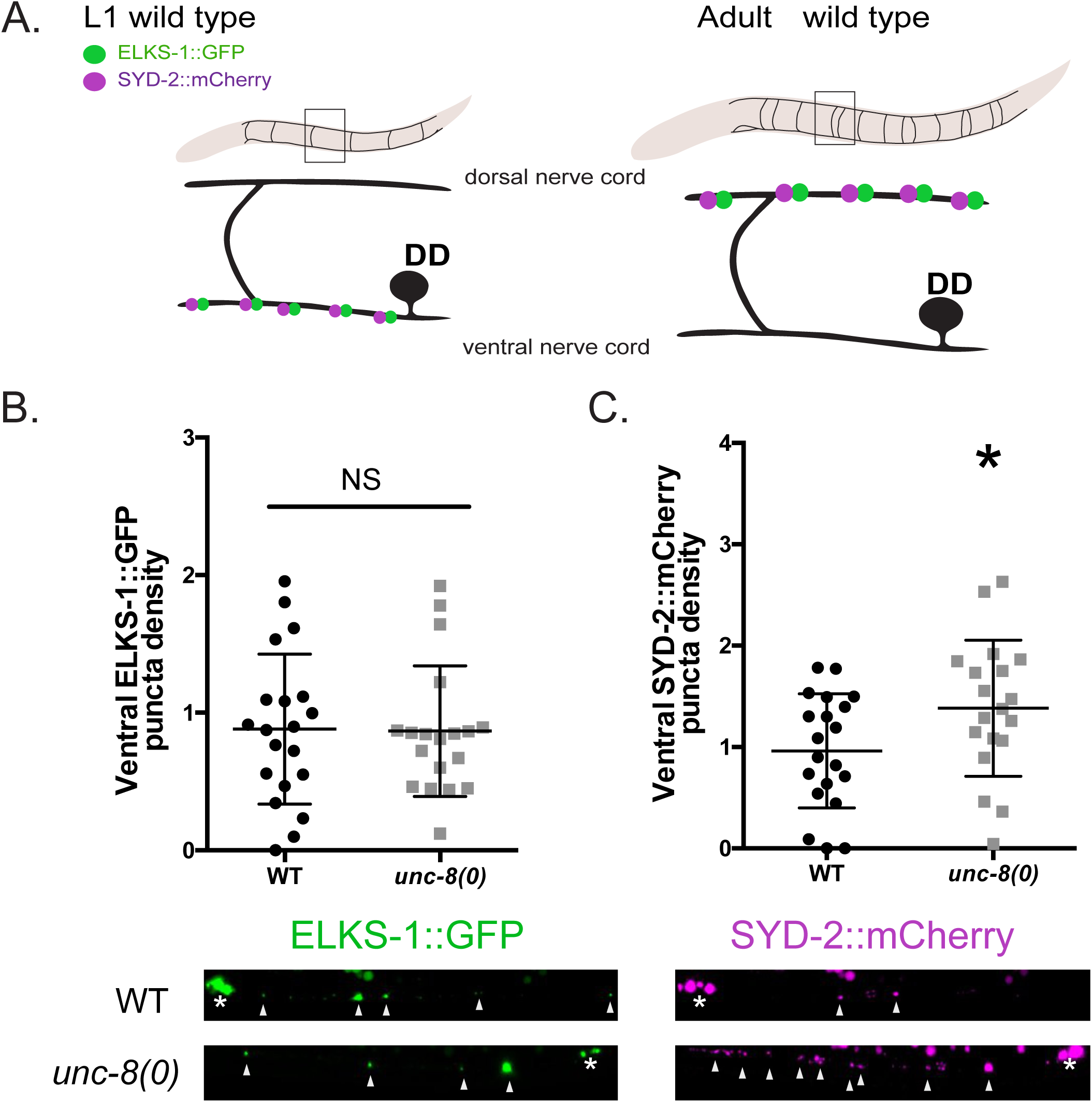
UNC-8 is required for the removal of SYD-2 but not ELKS-1 in remodeling DD motor neurons. **A.** Graphical representation of presynaptic ELKS-1::GFP (green) and SYD-2::mCherry (magenta) remodeling in wild-type DD motor neurons. Both presynaptic proteins are removed from DD synapses in the L1 larval ventral nerve cord (left) and relocated to DD synapses in the dorsal nerve cord as depicted in an adult (right). **B.** The density of ventral ELKS-1::GFP puncta (arrowheads) in wild-type (WT) (0.88 ± 0.5) and *unc-8* mutant worms (0.86 ± 0.4) are not significantly different at the young adult stage. Data are Mean ± SD. N > 19, unpaired T-test, p = 0.46. puncta density = puncta/10 μm. **C.** Ventral SYD-2::mCherry puncta (arrowheads) are retained in *unc-8* mutant worms (1.38 ± 0.7) compared to wildtype (WT) (0.96 ± 0.6) at the young adult stage. Data are Mean ± SD. N > 18, unpaired T-test, * p = 0.02. puncta density = puncta/10 μm. Scale bar = 10 μm.

**Figure 7.**
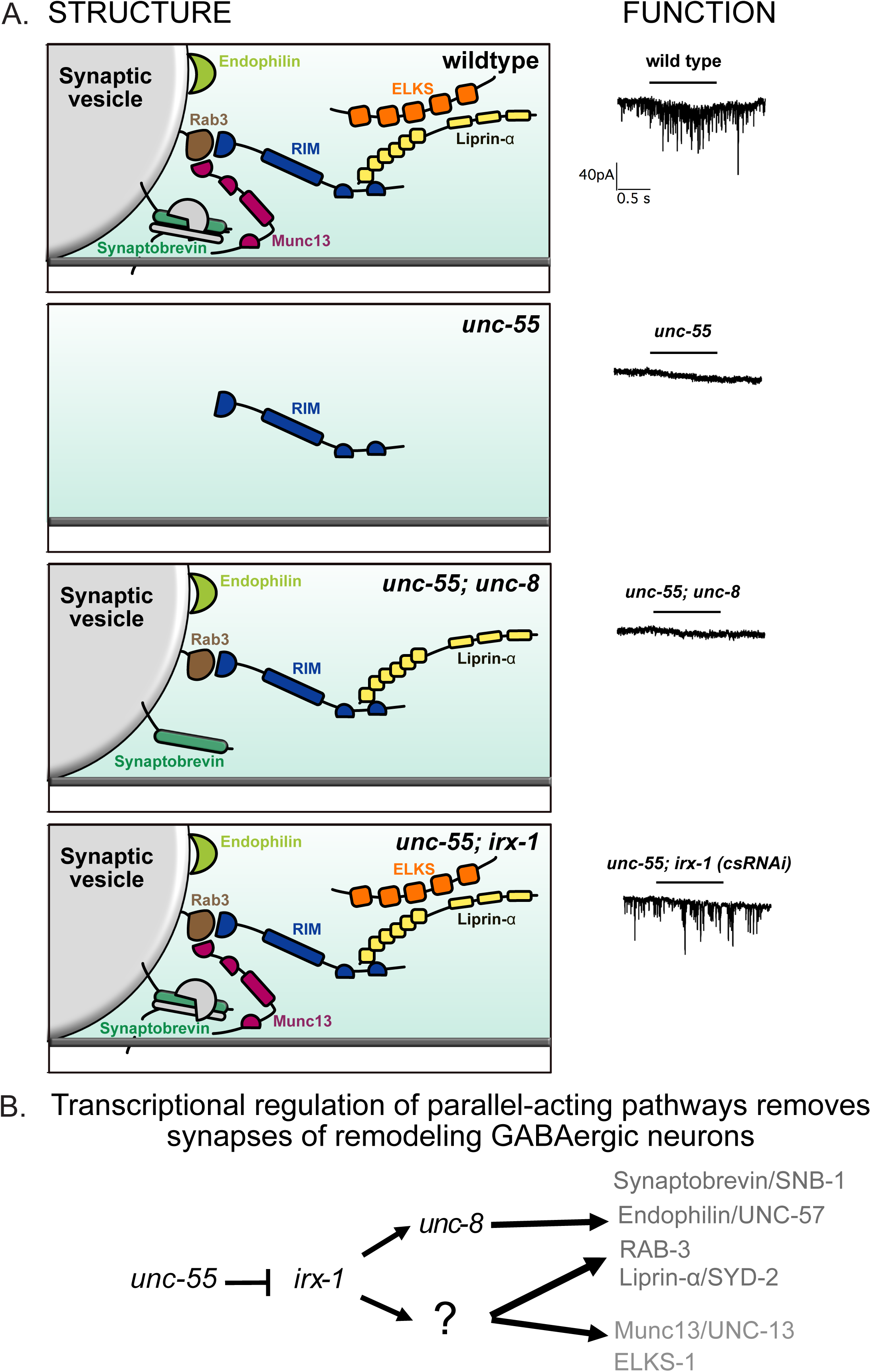
Parallel-acting pathways dismantle the presynaptic apparatus in remodeling GABAergic neurons. **A.** Structure and function of remodeling GABAergic synapses. wildtype: (Left) Key presynaptic proteins, Synaptobrevin, Rab3, RIM, Munc13, ELKS and liprin-*α* mediate synaptic vesicle fusion and spontaneous neurotransmitter release (Right) from ventral GABAergic synapses. *unc-55*: (Left) All of these presynaptic markers except RIM are removed from the ventral nerve cord in *unc-55* mutants in which both DD and VD GABAergic synapses are relocated to the dorsal nerve cord. (Right) As a result, *unc-55* mutants lack functional ventral GABAergic synapses. *unc-55; unc-8*: (Left) Genetic ablation of *unc-8* in *unc-55; unc-8* mutants prevents the removal of Synaptobrevin, Rab3, liprin-*α* and endophilin. Synaptic vesicles are docked by Rab3 and RIM near active zones, but do not fuse due to the absence of ELKS and Munc13 (Right). *unc-55; irx-1*: (Left) RNAi knockdown of the transcription factor gene *irx-1* blocks removal of Synaptobrevin, Rab3, liprin-*α*, endophilin, Munc13 and ELKS from ventral GABAergic synapses in *unc-55* mutants and results in the restoration of spontaneous ventral GABA release (Right). **B.** Transcriptional regulation of parallel-acting pathways for presynaptic disassembly. The COUP/TF transcription factor UNC-55 blocks expression of Iroquois/IRX-1 in VD GABAergic neurons to prevent presynaptic remodeling. In *unc-55* mutants, IRX-1 activates expression of the UNC-8/DEG/ENaC channel which then drives the removal of Synaptobrevin/SNB-1, Endophilin/UNC-57, RAB-3 and liprin-*α*/SYD-2. IRX-1 also activates additional downstream effectors (?) that promote the removal of ELKS-1 and Munc13/UNC-13 in addition to Synaptobrevin/SNB-1, Endophilin/UNC-57, RAB-3 and liprin-*α* via *unc-8*-independent mechanisms.

As an additional test of this idea, we expressed ELKS-1::GFP and SYD-2::mCherry in DD neurons for simultaneous dual color imaging. This experiment confirmed that more SYD-2::mCherry puncta are retained in the ventral nerve cords of adult *unc-8* mutants than wildtype, thus confirming that *unc-8* normally promotes the removal of SYD-2 from ventral DD synapses in the native remodeling program (Figure 6). In contrast, few ventral DD ELKS-1::GFP are retained in either wild-type or *unc-8* mutant adults, an observation consistent with our finding above (Figure 5B) that UNC-8 function is not required for the removal of ELKS-1 from GABAergic synapses that remodel in *unc-55* mutants. Overall, our results demonstrate that subsets of active zone proteins are targeted for removal by separate mechanisms that disassemble the presynaptic apparatus in remodeling GABAergic neurons (Figure 7).

## DISCUSSION

### Presynaptic disassembly in remodeling in GABAergic neurons

Synaptic plasticity is a key dynamic feature of the nervous system as neurons actively assemble new synapses while dismantling others (Chen et al. 2011; Nishiyama et al. 2007; De Paola et al. 2006). In contrast to synaptic assembly about which much is known, the molecular mechanisms that drive synaptic elimination are relatively unexplored (T. Südhof 2018; T. C. Südhof 2017). In this study, we investigated a developmentally-regulated mechanism of presynaptic disassembly in *C. elegans* (White, Albertson, and Anness 1978). Our findings revealed parallel-acting pathways that selectively remove different components of the presynaptic active zone in remodeling GABAergic synapses (Figure 7). We have shown that the conserved transcription factor Iroquois/IRX-1 drives expression of the DEG/ENaC channel subunit UNC-8 (Figure 2) to remove the presynaptic proteins Synaptobrevin/SNB-, liprin-*α*/SYD-2, Endophilin/UNC-57 and RAB-3 (Figure 3A-C). IRX-1 regulates an additional parallel-acting pathway that also removes these presynaptic components (Figure 3D-G). In addition, Iroquois/IRX-1 promotes the disassembly of ELKS-1 and Munc13/UNC-13 in a separate pathway that does not require UNC-8 activity (Figure 4, 5, 6 and 7). Together, our findings support the conclusion that synaptic disassembly can be transcriptionally-regulated and involve molecularly distinct mechanisms that differentially eliminate selected subsets of presynaptic proteins.

### Activity-dependent active zone remodeling

The active zone region of the presynaptic terminal mediates synaptic vesicle (SV) fusion for neurotransmitter release (T. C. Südhof 2012). This active zone function is defined by a core group of components including Voltage-gated Ca^++^ Channels (VGCCs), ELKS, Munc13/UNC-13, liprin-*α*/SYD-2, SYD-1, RIM/UNC-10, and Rim Binding Protein (RBP) (T. C. Südhof 2012). Notably, the composition and size of the SV release machinery can by modulated by synaptic activity. For example, additional copies of specific active zone proteins (i.e. ELKS, RBP, VGCCs and Munc13) are incorporated into the presynaptic zones of *Drosophila* neuromuscular junctions (NMJs) in a homeostatic mechanism that elevates neurotransmitter release to compensate for reduced postsynaptic sensitivity (Böhme et al. 2019; Gratz et al. 2019). Elevated activity in *Drosophila* photoreceptors can also have the opposite effect of selectively removing a subset of these presynaptic proteins (liprin-*α*, RIM and RBP) while leaving others intact (VGCCs and SYD-1) (Sugei et al. 2015). Our findings point to a parallel effect in remodeling GABAergic neurons in *C. elegans* in which neuronal activity promotes the elimination of selected presynaptic components. In previous work, we determined that the DEG/ENaC channel UNC-8 functions in an activity-dependent pathway that dismantles the presynaptic active zone. Genetic results, for example, show that UNC-8 acts in a common pathway with the VGCC, UNC-2 (Miller-Fleming et al. 2016). Thus, we propose here that UNC-8-driven removal of Synaptobrevin/SNB-1, liprin-*α*/SYD-2, Endophilin/UNC-57 and RAB-3 also depends on GABA neuron activity and cytoplasmic calcium whereas synaptic elimination of Munc13 and ELKS, which does not require UNC-8, is selectively regulated in a separate pathway driven by the transcription factor Iroquois/IRX-1 (Figure 7). Future studies are needed to define the downstream IRX-1 effectors that promote Munc13 and ELKS elimination. Because Iroquois/IRX-1 expression and synaptic remodeling in GABAergic neurons are blocked by the transcription factor UNC-55 (He et al. 2015; Petersen et al. 2011; Zhou and Walthall 1998), molecular regulators of Munc13 and ELKS elimination may be included in previously defined data sets of UNC-55-regulated genes (Petersen et al. 2011; Yu et al. 2017).

## ACKOWLEDGEMENTS

We thank K. Shen for the ELKS-1::GFP and SYD-2::mCherry markers used in Figure 6, J. Kaplan, and J. Dittman for additional reagents and M. Kittelman and M. Zhen for advice on scoring GABA synapses by electron microscopy. Some strains used in this study were provided by the CGC, which is funded by the NIH Office of Research Infrastructure Programs (P40 OD010440) This work used instruments in the Electron Microscopy Core of the UIC Research Resources Center; the BioCryo facility of Northwestern University’s NUANCE Center, which has received support from the Soft and Hybrid Nanotechnology Experimental (SHyNE) Resource (NSF ECCS-1542205); the MRSEC program (NSF DMR-1720139) at the Materials Research Center; the International Institute for Nanotechnology (IIN); and the State of Illinois, through the IIN; and CryoCluster equipment, which has received support from the MRI program (NSF DMR-1229693). Imaging experiments were performed in part in the Vanderbilt Cell Imaging Shared Resource (supported by NIH grants CA68485, DK20593, DK58404, DK59637 and EY08126). This work was supported by NIH grants R01NS081259, R01NS106951(DMM) and predoctoral fellowships from the NIH to TWFM (1F31NS084732), NSF to SP (DGE:1445197) and AHA to SP (19PRE34380582) and ACC (18PRE33960581).

## DECLARATION OF INTERESTS

The authors declare no financial interests.

## METHODS

### Strains and Genetics

*C. elegans* strains were cultured at either 20° C or 23° C as previously described on standard nematode growth medium seeded with OP50 (Brenner 1974). The mutant alleles and strains used in this study are outlined in **Tables 1** and **2**.

**Table 1.**
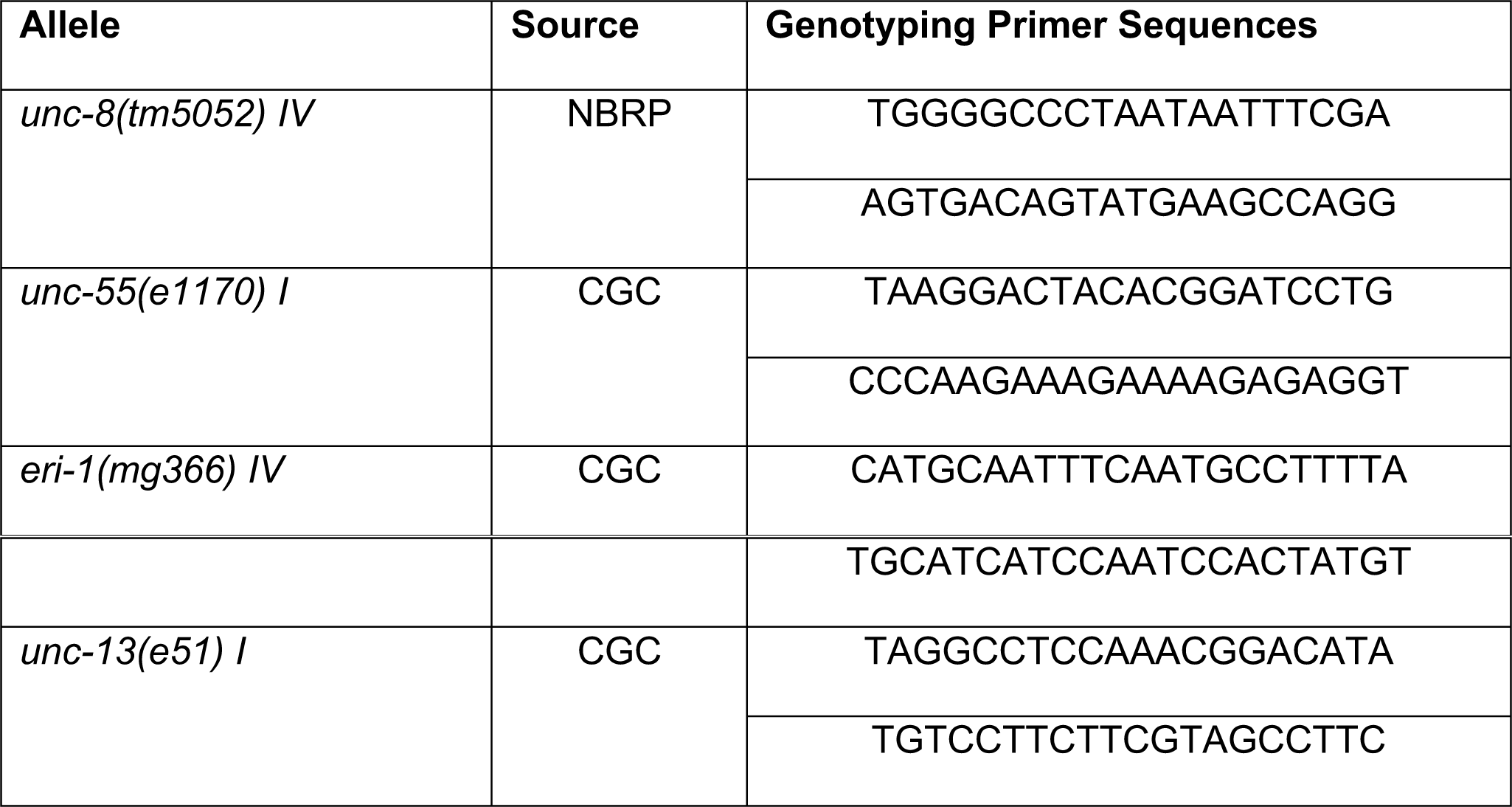
Mutant alleles and genotyping primers used in this study.

**Table 2.**
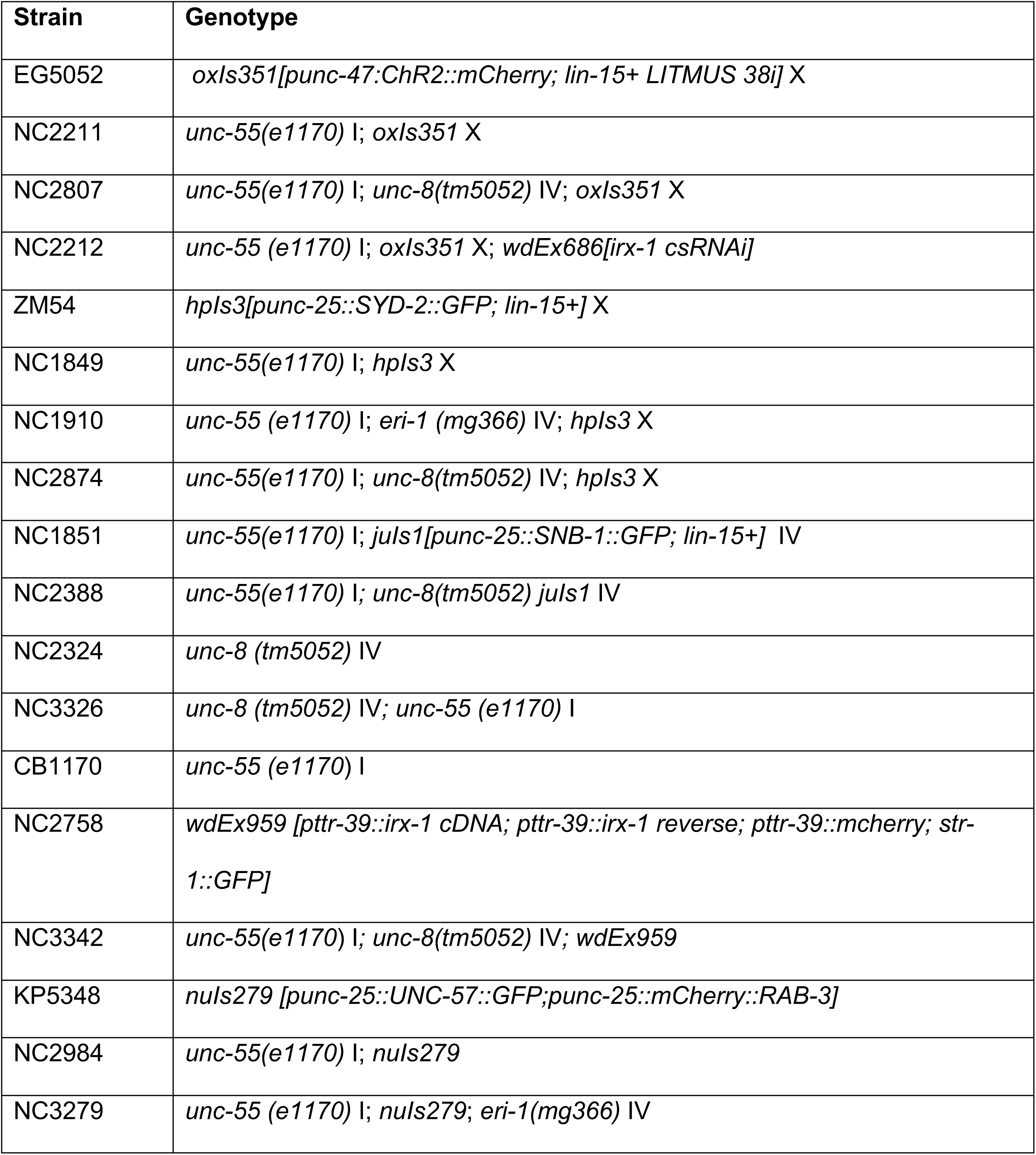
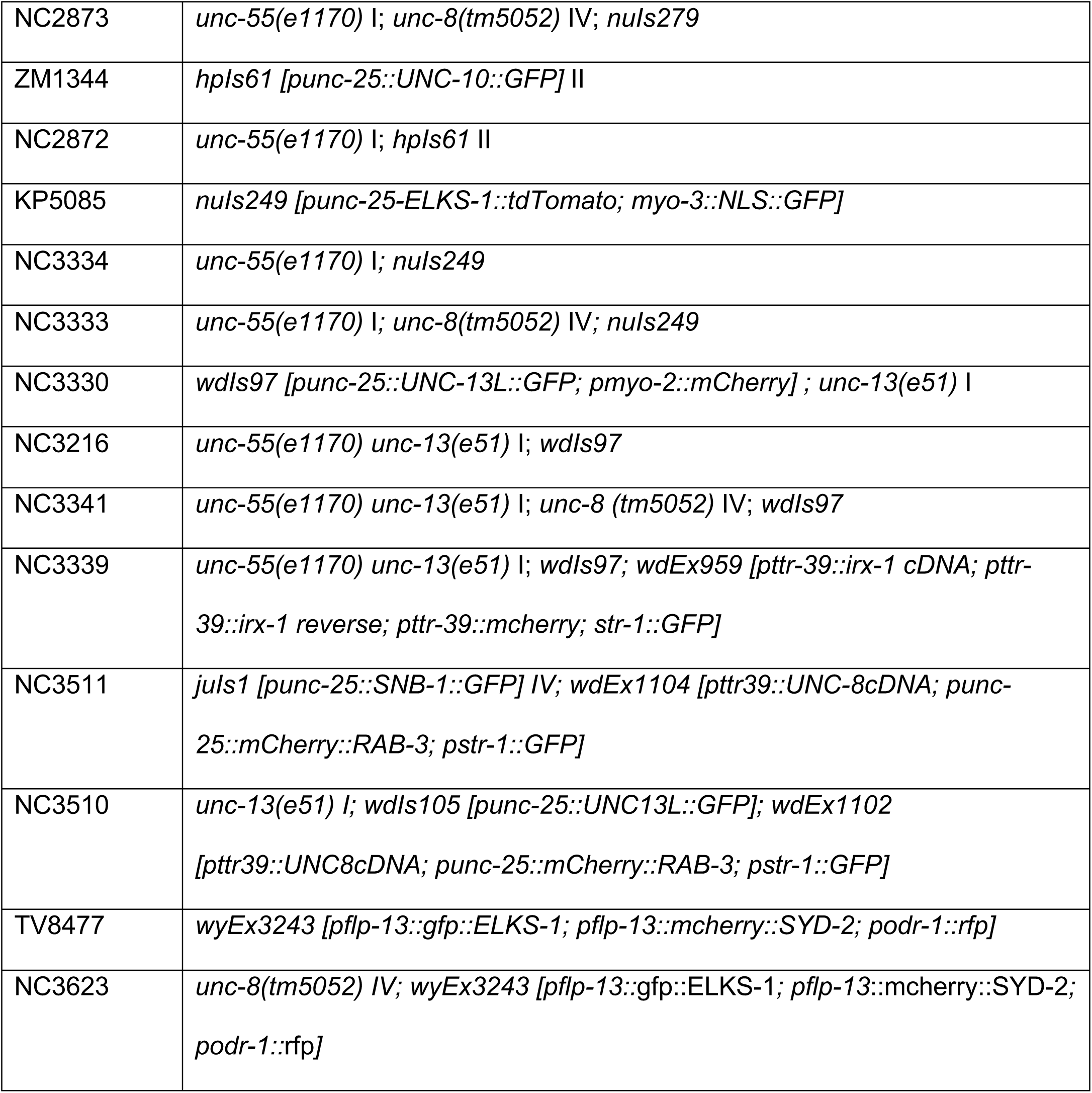
Strains used in this study.

### Microscopy

#### Confocal Microscopy

Larval or young adult animals were immobilized on 2% agarose pads with 15mM levamisole as previously described. Z-stack images (Figure 3A-C, 5, 5S1B-C) were collected on a Leica TCS SP5 confocal microscope using a 63X oil objective (0.5 μm/step), spanning the focal depth of the ventral nerve cord GABA neurons and synapses. Leica Application Suite Advanced Fluorescence (LAS-AF) software was used to generate maximum intensity projections. Ventral nerve cord images were straightened using an ImageJ plug-in. Z-stack images (Figure 3D-F, 4S3, 5S1A, 5S2 and 5S3) were acquired with Nikon confocal A1R using 40X/1.3 and 60X/1.4 N.A. oil objective (0.5 μm/step).

#### Image Analysis

Synapse density counts (Figures 3A-C and 5) were collected by tracing segments of the ventral nerve cord using the segmented line tool in ImageJ. Distance in micrometers and gray value plot traces were used to count the number of peaks (synapses) that occur over the specified distance. Synapses were defined as fluorescent peaks that reached a threshold of 25 arbitrary units of fluorescence intensity.

NIS Elements 5.2 software was used to produce Figures 2, 3D-F, 4S3 and 5S1. Synaptic density for each marker (UNC-10::GFP, SYD-2::GFP, RAB-3::mCherry, UNC-57::GFP and SNB-1::GFP) was calculated using the General Analysis tool. First, images were preprocessed to subtract background using Rolling Ball Correction. Then, the intensity threshold was defined for each marker and binary objects were filtered by size and circularity. Each object along the nerve cord was considered a synaptic punctum. Density was defined as the number of puncta per 10 μm of dendrite.

FIJI was used to produce Figures 6 and 5S3. Line scans were drawn on the ventral cord anterior to the VD cell body of interest. Background fluorescence was obtained from a line scan from an adjacent region interior to the ventral cord. Average intensity was reported for both line scans and subtracted (Figure 5S3). Puncta density was counted in the GFP and mCherry channel by a scorer blinded to the genotype (Figure 6).

### Single molecule mRNA Fluorescence In Situ Hybridization (smFISH)

smFISH was performed with custom *unc-8* probes linked to Quasar® 670 (Biosearch Technologies). Synchronized larvae (from either late L1 or early L3 stage) were collected by washing plates with M9, fixed in 4% paraformaldehyde in 1X PBS for 45 min and permeabilized in 70% ethanol for 48 h. Hybridization followed the manufacturer’s instructions (http://www.biosearchtech.com/stellarisprotocols) and was performed at 37°C for 16h in Stellaris RNA FISH hybridization buffer (Biosearch Technologies Cat# SMF-HB1-10) containing *unc-8* probe at 1:100. For *irx-1* cell specific RNAi (csRNAi) experiments, all DD motor neurons were marked with *Punc-47*::GFP (*oxIs12*) and specific DDs expressing the *irx-1(csRNAi)* constructs (*pttr-39::irx-1* sense, *pttr-39::irx-1* antisense) were co-labeled with *Punc-25::*mCherry to distinguish them from DD neurons that did not express the *irx-1(csRNAi)* transgenic array. For *irx-1(oe)* (*irx-1* over-expression) experiments, GFP-tagged IRX-1 was expressed with the *ttr-39* promotor (*pttr-39::IRX-1::*GFP). In this setup, VDs and DDs were marked with *Punc-47::mCherry* (*wpIs39*) and individual DDs or VDs expressing *irx-1(oe)* were detected by expression of nuclear-localized IRX-1::GFP (Petersen et al. 2011). In all cases, cell nuclei were stained with DAPI. Z-stacks were collected in a Nikon spinning disk confocal microscope with optical filters for DAPI, Quasar® 670 and GFP using a 100X objective (NA=1.49) in 0.2 μm steps spanning the cell body and merged for quantification following 3D-deconvolution in Nikon elements. smFISH puncta were counted if they corresponded to circular fluorescent spots, exceeded the Quasar® 670 background signal and were located within either a GFP-labeled or mCherry-marked DD/VD cell body. At least 30 worms were scored for each group and the Mann-Whitney test used to determine significance (n >45 neurons). As a positive control, *unc-8* smFISH staining was noted in adjacent DA and DB ventral-cord neurons for all samples to confirm successful hybridization.

### Electron Microscopy

Young adult hermaphrodites of each strain were prepared for high-pressure freeze (HPF) fixation as described (Miller-Fleming et al. 2016; Rostaing et al. 2004). 10–15 animals were loaded into a specimen chamber filled with *E. coli.* The specimens were frozen rapidly in a high-pressure freezer (Leica HPM100) at -180°C and high pressure. Freeze substitution was performed on frozen samples in a Reichert AFS machine (Leica, Oberkochen, Germany) with 0.1% tannic acid and 2% OsO4 in anhydrous acetone. The temperature was kept at -90°C for 107 h, increased at 5°C/h to -20°C, and kept at -20°C for 14h. The temperature was then increased by 10°C/h to 20°C. Fixed specimens were embedded in Epon resin after infiltration in 50% Epon/acetone for 4h, 90% Epon/acetone for 18h, and 100% Epon for 5 hours. Embedded samples were incubated for 48h at 65°C. All specimens were prepared in the same fixation procedure and labeled with anonymous tags so that the examiner was blinded to genotype. Ultrathin (40 nm) serial sections were cut using an Ultracut 6 (Leica) and collected on formvar-covered, carbon-coated copper grids (EMS, FCF2010-Cu). Grids were counterstained in 2% aqueous uranyl acetate for 4 min, followed by Reynolds lead citrate for 2 min. Images were obtained on a Jeol JEM-1220 (Tokyo, Japan) transmission electron microscope operating at 80 kV. Micrographs were collected using a Gatan digital camera (Pleasanton, CA) at a magnification of 100k. Images were quantified using NIH ImageJ software. Dorsal and ventral cords were distinguished by size and morphology. GABAergic synapses were identified by previously established criteria, including position in the cord as well as the morphology of the synapse. GABAergic synapses are larger than their cholinergic motor neuron counterparts, and the active zones in these synapses form a direct, perpendicular angle with muscle arms. In contrast, the presynaptic density in cholinergic synapses orient at an acute angle to the muscle, generally 30-45° and are often dyadic. Some images were collected at 30k to aid in identifying synaptic identity based on terminal position in the cord. Two colleagues with expertise in EM reconstruction of the *C. elegans* ventral nerve cord independently reviewed synapse images from each strain to verify identification. Each profile represents an image of a 40 nm section. A synapse was defined as a set of serial sections containing a presynaptic density with flanking sections from both sides without presynaptic densities. Synaptic vesicles were identified as spherical, light gray structures with an average diameter of ∼30 nm. Synaptic vesicles were considered docked if they touched the membrane. Three to five animals were imaged for each genotype. Numbers of profiles for each genotype were (# analyzed / # imaged): wild type = 80/1330, *unc-55; unc-8* = 37/745, *unc-55;irx-1(csRNAi)* = 54/613 for ventral GABAergic synapse evaluation.

### Electrophysiology

The *C. elegans* dissection and electrophysiological methods were as previously described (Miller-Fleming et al. 2016; Richmond and Jorgensen 1999). Animals were immobilized along the dorsal axis with Histoacryl Blue glue, and a lateral cuticle incision was made with a hand-held glass needle, exposing ventral medial body wall muscles. Muscle recordings were obtained in the whole-cell voltage-clamp mode using an EPC-10 patch-clamp amplifier and digitized at 1 kHz. The extracellular solution consisted of 150 mM NaCl, 5 mM KCl, 5 mM CaCl2, 4 mM MgCl2, 10 mM glucose, 5 mM sucrose, and 15 mM HEPES (pH 7.3, ∼340 mOsm). The intracellular solution consisted of 120 mM KCl, 4 mM KOH, 4 mM MgCl2, 5 mM (N-tris[Hydroxymethyl] methyl-2-aminoethane-sulfonic acid), 0.25 mM CaCl2, 4 mM Na2ATP, 36 mM sucrose, and 5 mM EGTA (pH 7.2, ∼315 mOsm). GABAergic mininIPSCs and hyperosmotic responses were acquired at a holding potential of -60 mV by pressure-ejecting extracellular saline containing an additional 500 mOsm of sucrose. 10 mM d-tubocurare was added to both the extracellular solution and the pressure ejection pipette to block cholinergic hyperosmotic currents. Data were acquired using Pulse software (HEKA, Southboro, Massachusetts, United States) on a Dell computer. Subsequent analysis and graphing were performed using Pulsefit (HEKA), Mini analysis (Synaptosoft Inc., Decatur, Georgia, United States) and Igor Pro (Wavemetrics, Lake Oswego, Oregon, United States).

### Molecular Biology

#### Generation of the punc-25::UNC-13L::GFP transgenic line

We used the In-Fusion cloning kit (Takara) to amplify the cDNA of the long isoform of UNC-13 (UNC-13L) from a plasmid provided by J. Kaplan (pTWM88). This fragment was ligated into a vector containing the *punc-25* GABA promoter and a C-terminal GFP tag. The resulting plasmid, pTWM90, was injected into *unc-13 (e51)* null mutants at 25 ng/µl with the co-injection marker *pmyo-2::mCherry* (2 ng/ µl). This transgenic array was integrated by x-ray irradiation and outcrossed for three generations to generate stable transgenic lines for analysis.

### Feeding RNA Interference Experiments

Bacteria producing either double-stranded *irx-1* RNA or containing the RNAi empty vector were seeded on NGM plates and stored at 4°C for up to 1 week. Four L4 *unc-55, unc-55; eri-1,* or *unc-55; unc-8* animals were grown on each single RNAi plate at 23°C until progeny reached the L4 stage. Progeny were picked to fresh RNAi plates and the ventral synapses were quantified.

### Movement Assays

Animals were first tapped on the tail to ensure that they were capable of forward locomotion, then tapped on the head to assess ability to execute backward locomotion. Animals were binned into the following categories: “unc” (uncoordinated: coil ventrally immediately upon tapping), “initiate backing” (initiate backwards movement but stop), and “wild-type” (sustain backward locomotion with at least two body bends). In Figure 4S1, the “wild-type” and “initiate backing” categories were grouped into a single “initiate backing” category.

### IRX-1 cell-specific RNAi

The *irx-1(*csRNAi) array (*wdEx959*) was outcrossed from NC2975 (He et al. 2015). The new outcrossed line NC2758 was crossed with the *punc-25*::UNC-13L::GFP transgene (Figure 5A).

## SUPPLEMENTARY IMAGES

**FIGURE 4 – SUPPLEMENT 1.**
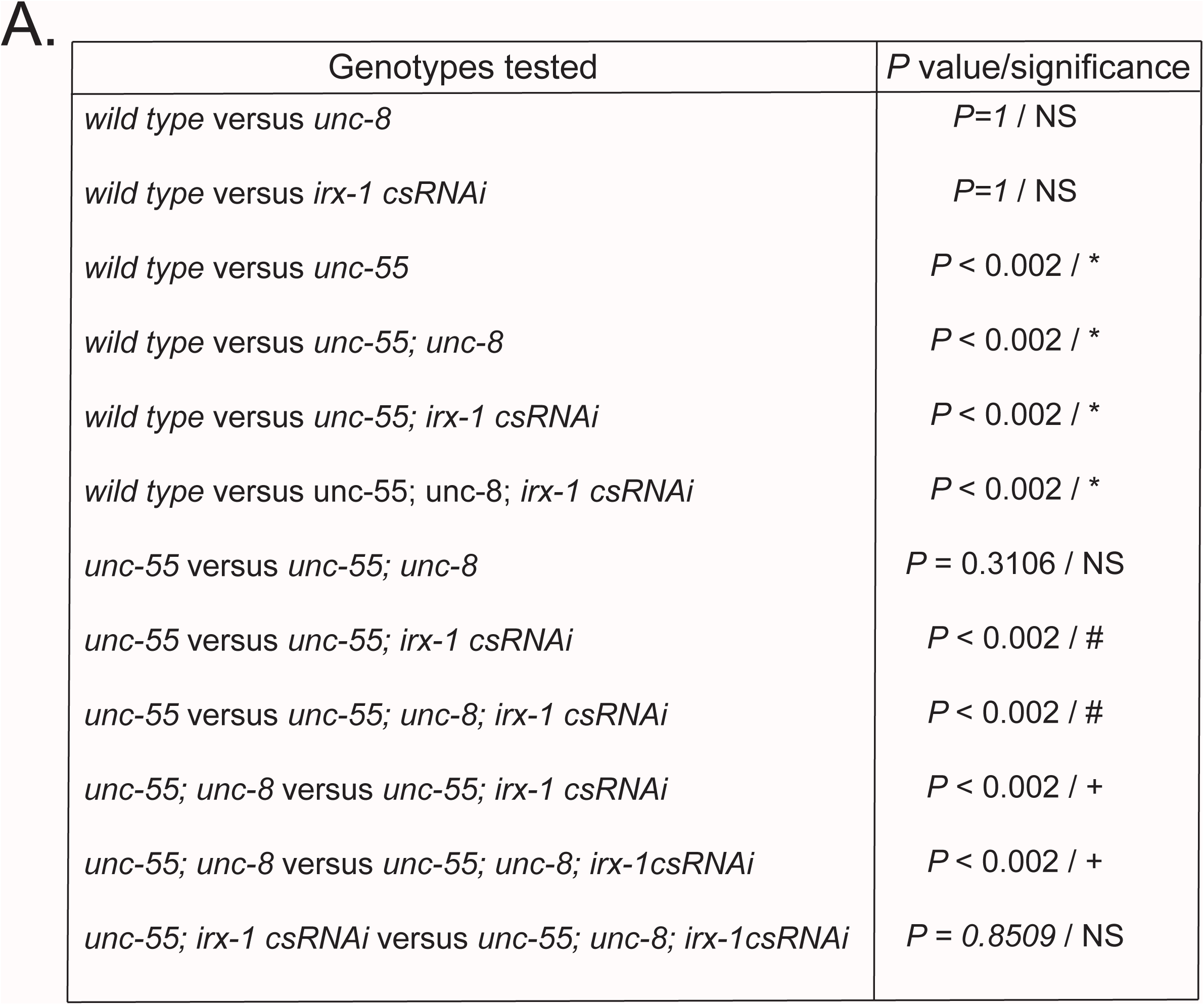
**A.** Table that compiles all genotypes used in the locomotion assay. Fisher’s exact test was used for pairwise comparisons.

**FIGURE 4 – SUPPLEMENT 2.**
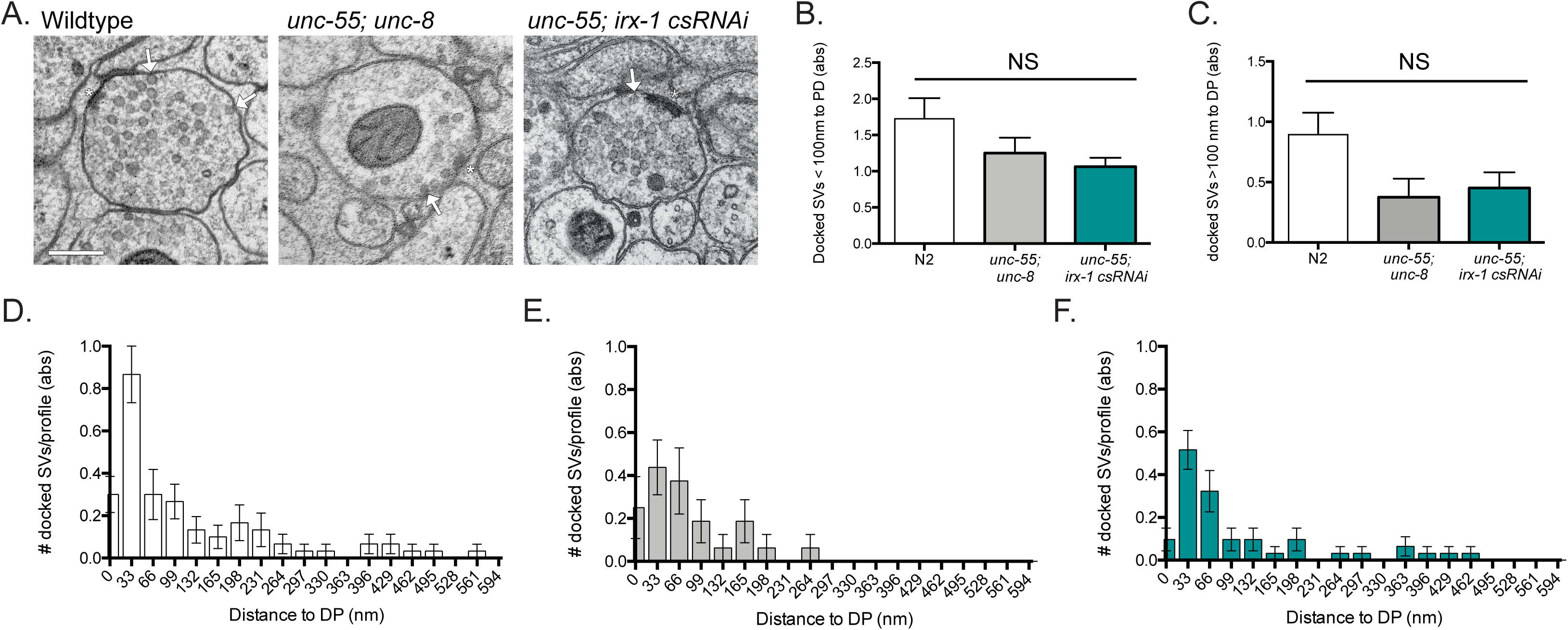
**A.** Number of docked synaptic vesicles < 100 nm of the dense projection (**B**) or > 100 nm from the dense projection (**C**) in each profile of ventral GABAergic synapses, where distance to the dense projection is measured along the length of the plasma membrane in a single section. Across genotypes, more docked vesicles are observed within 100 nm of the dense projection than distal to this region (see **D-F**). An apparent reduction in docked SVs in the mutant genotypes at distances of > 100 nm is not significant (p = 0.1312). Data were analyzed by Kruskal-Wallis. **D-F.** Distances between each docked synaptic vesicle and the dense projection were measured and sorted into 33 nm bins for wild type (**D**), *unc-55; unc-8* (**E**), and *unc-55; irx-1(csRNAi)* (**F**) animals. Docked synaptic vesicles are localized more closely to the dense projection and appear less frequently at greater distances. In *unc-55; unc-8* mutant animals, no docked SVs were observed > 264 nm from the dense projection (**E**).

**FIGURE 4 – SUPPLEMENT 3.**
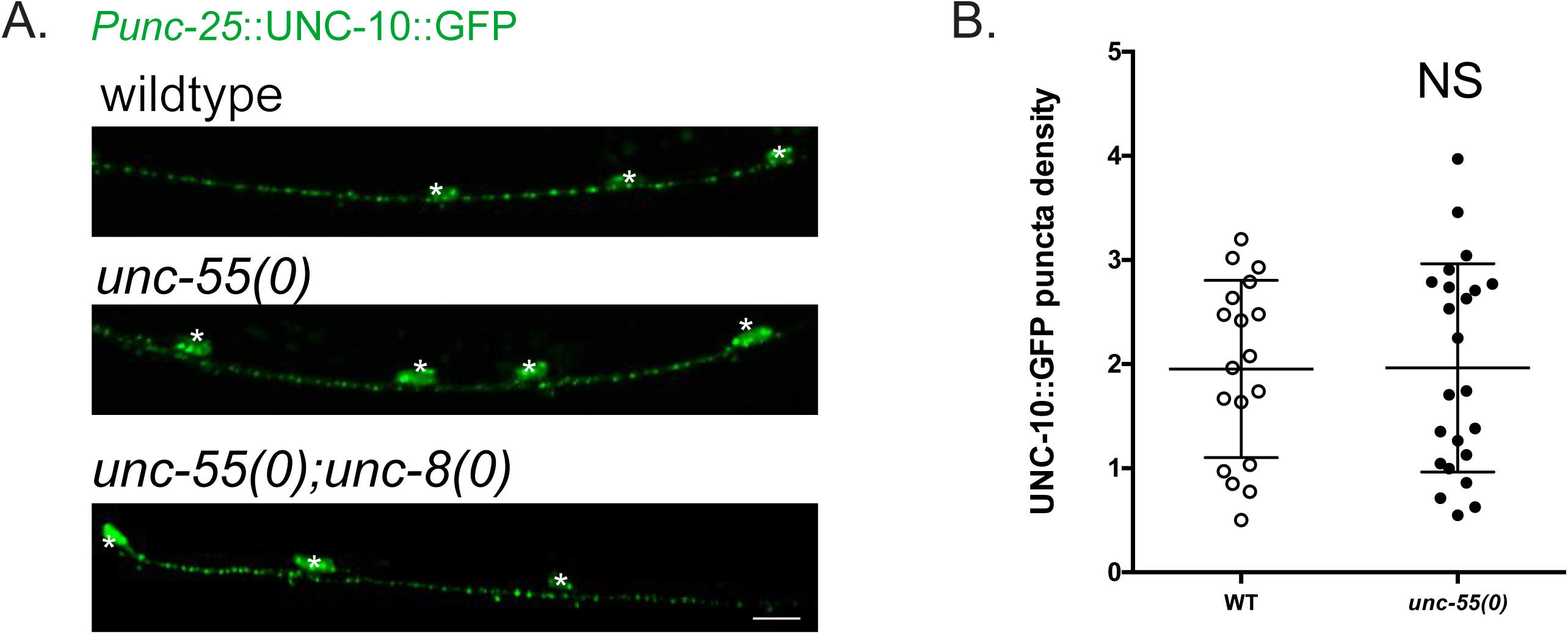
UNC-10::GFP persists in the ventral cord of *unc-55* mutant animals. **A.** Representative images of UNC-10::GFP in the ventral cord of wild-type (WT), *unc-55* mutants and *unc-55; unc-8* double mutant worms. Scale bar = 5µm. **B.** Quantification of UNC-10::GFP puncta in the ventral cord. UNC-10::GFP puncta density does not differ between wild-type (WT) (1.95 ± 0.9) vs *unc-55* mutant worms (1.96 ± 1.0). Data are Mean ± SD. N > 17. n.s. = not significant. p = 0.486.

**FIGURE 5 – SUPPLEMENT 1.**
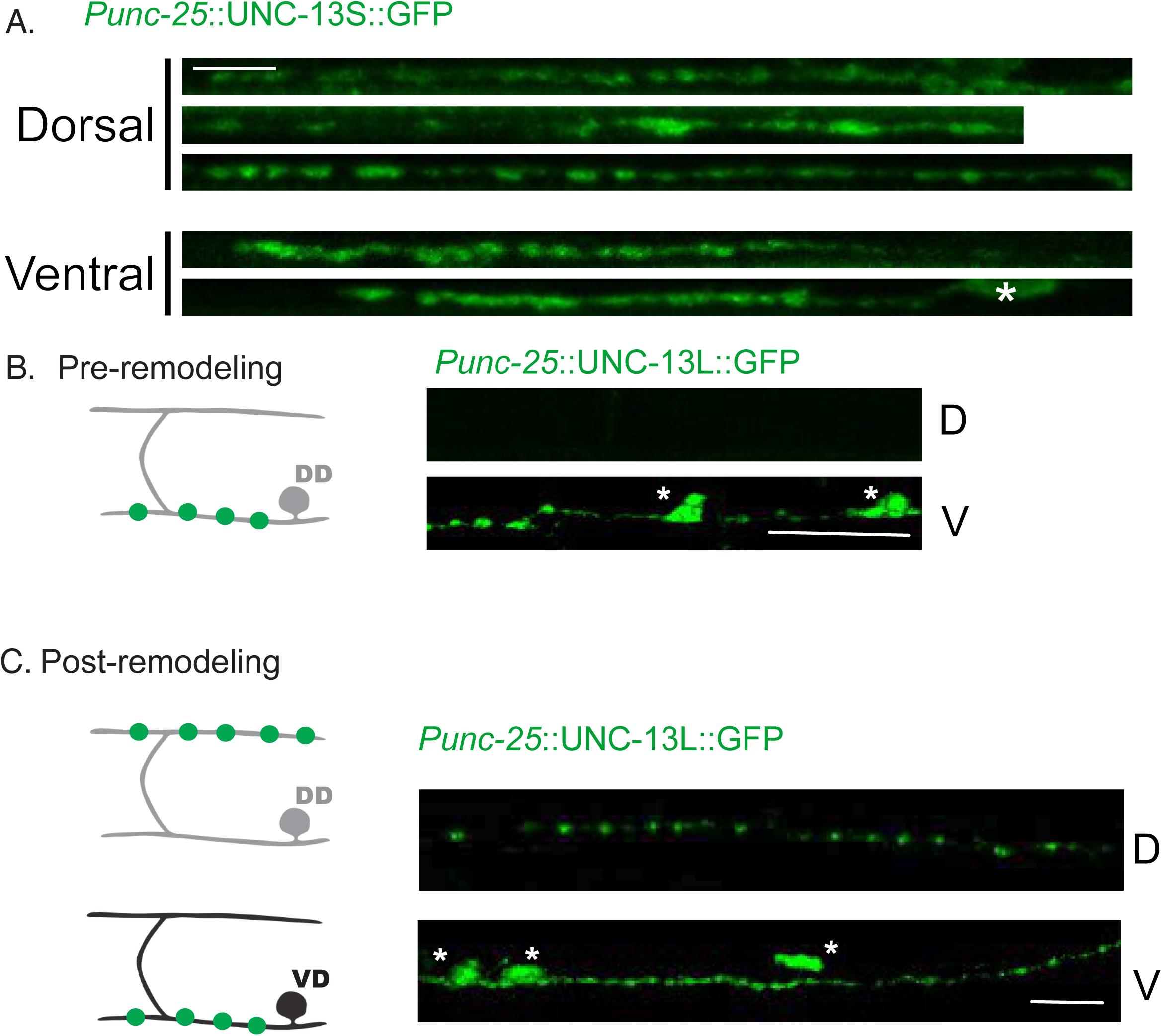
UNC-13L::GFP remodels in DD neurons and is localized to GABAergic synapses in VD neurons. **A.** Fluorescently-tagged UNC-13S *(punc-25::UNC-13S::GFP)* is diffused at presynaptic boutons of GABAergic motor neurons. Scale bar = 5µm. **B.** Fluorescently-tagged UNC-13L *(punc-25::UNC-13L::GFP)* is localized to the ventral (V) but not dorsal (D) nerve cords before remodeling (in L1 animals). Scale bar = 5µm. **C.** UNC-13L::GFP is visible in both the dorsal (D) and ventral (V) nerve cords after remodeling at the L4 stage. Scale bar = 5µm.

**FIGURE 5 – SUPPLEMENT 2.**
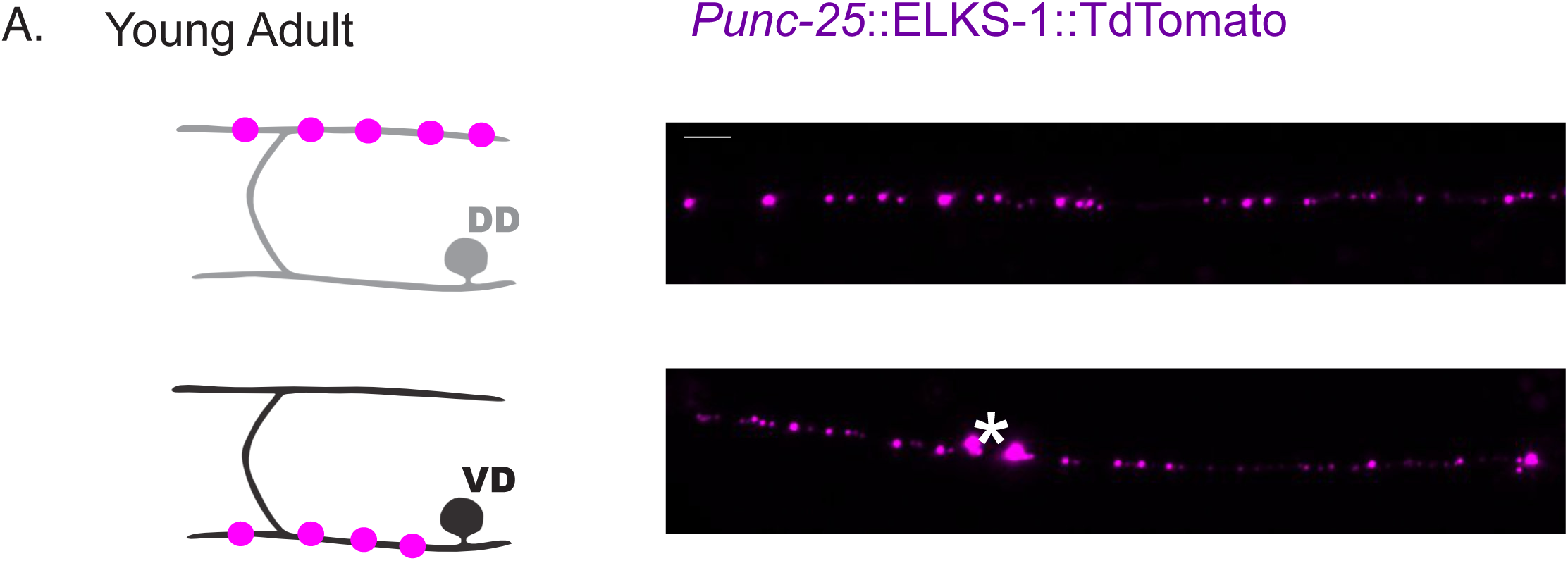
ELKS-1::TdTomato localizes to presynaptic terminals of DD neurons and VD neurons **A.** ELKS-1::TdTomato is visible in both the dorsal (D) and ventral (V) nerve cords after remodeling at the L4 stage. Scale bar = 5µm.

**FIGURE 5 – SUPPLEMENT 3.**
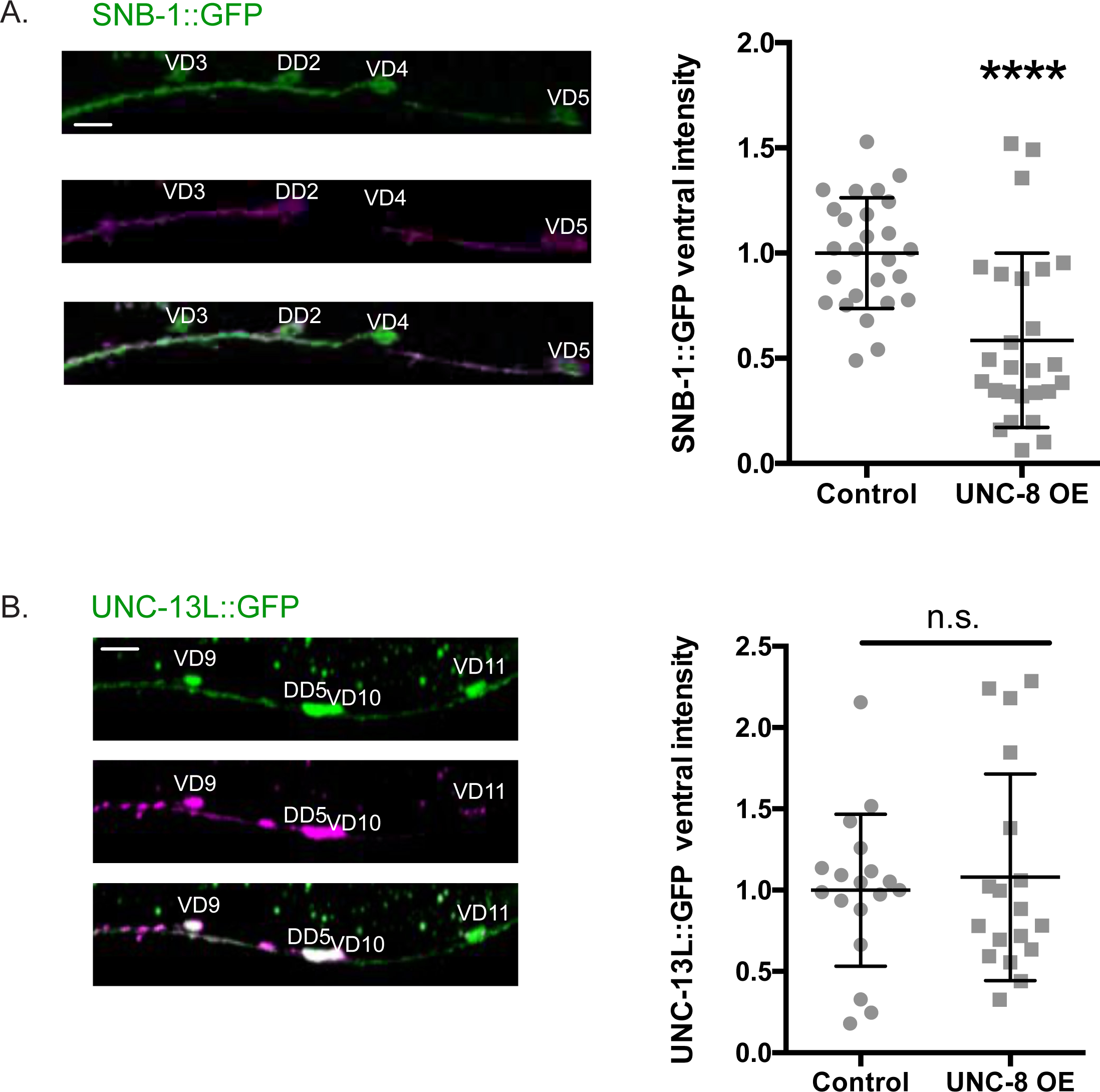
UNC-8 is sufficient to remove SNB-1 but not UNC-13L from ventral VD neuron GABAergic synapses. **A.** Forced expression of UNC-8 in VD neurons (magenta) (1.00 ± 0.3) reduces SNB-1::GFP (green) levels in GABAergic terminals compared to control VD neurons that do not express UNC-8 (0.58 ± 0.4). N = 24 animals. **** p < 0.0001. Unpaired T-test. **B.** Forced expression of UNC-8 in VD neurons (magenta) (1.00 ± 0.5) is not sufficient to reduce UNC-13 levels in ventral VD neuron GABAergic terminals compared to control VD neurons that do not express UNC-8 (1.08 ± 0.6). Data are Mean ± SD. N = 18. NS = not significant. p = 0.546. Unpaired T-test.

